# Mathematical modeling indicates that regulatory inhibition of CD8^+^ T cell cytotoxicity can limit efficacy of IL-15 immunotherapy in cases of high pre-treatment SIV viral load

**DOI:** 10.1101/2023.01.09.523305

**Authors:** Jonathan W. Cody, Amy L. Ellis-Connell, Shelby L. O’Connor, Elsje Pienaar

## Abstract

Immunotherapeutic cytokines can activate immune cells against cancers and chronic infections. N-803 is an IL-15 superagonist that expands CD8^+^ T cells and increases their cytotoxicity. N-803 also temporarily reduced viral load in a limited subset of non-human primates infected with simian immunodeficiency virus (SIV), a model of HIV. However, viral suppression has not been observed in all SIV cohorts and may depend on pre-treatment viral load and the corresponding effects on CD8^+^ T cells. Starting from an existing mechanistic mathematical model of N-803 immunotherapy of SIV, we develop a model that includes activation of SIV-specific and non-SIV-specific CD8^+^ T cells by antigen, inflammation, and N-803. Also included is a regulatory counter-response that inhibits CD8^+^ T cell proliferation and function, representing the effects of immune checkpoint molecules and immunosuppressive cells. We simultaneously calibrate the model to two separate SIV cohorts. The first cohort had low viral loads prior to treatment (≈3-4 log viral RNA copy equivalents (CEQ)/mL), and N-803 treatment transiently suppressed viral load. The second had higher pre-treatment viral loads (≈5-7 log CEQ/mL) and saw no consistent virus suppression with N-803. The mathematical model can replicate the viral and CD8^+^ T cell dynamics of both cohorts based on different pre-treatment viral loads and different levels of regulatory inhibition of CD8^+^ T cells due to those viral loads (i.e. initial conditions of model). Our predictions are validated by additional data from these and other SIV cohorts. While both cohorts had high numbers of activated SIV-specific CD8^+^ T cells in simulations, viral suppression was precluded in the high viral load cohort due to elevated inhibition of cytotoxicity. Thus, we mathematically demonstrate how the pre-treatment viral load can influence immunotherapeutic efficacy, highlighting the in vivo conditions and combination therapies that could maximize efficacy and improve treatment outcomes.

**Author Summary:** Immunotherapy bolsters and redirects the immune system to fight chronic infections and cancers. However, the effectiveness of some immunotherapies may depend on the level of pre-treatment inflammation and the corresponding presence of regulatory cells and immune checkpoint molecules that normally function to prevent immune overreaction. Here, we consider two previously published cohorts of macaques who were given the immunotherapeutic N-803 to treat Simian Immunodeficiency Virus, an analog of Human Immunodeficiency Virus (HIV). One cohort had low viral loads before treatment, and N-803 temporarily suppressed viral loads. The second cohort had high viral loads that did not consistently decrease with N-803 treatment. In this work, we demonstrate with a mathematical model how these two distinct outcomes can arise due only to the different viral loads and the corresponding immune activation and regulatory response. In the model, we find that the key limiting factor is the direct inhibition of the cytotoxic action of immune cells by immune checkpoint molecules. This model indicates that simultaneous blockade of immune checkpoint molecules may be necessary for effective application of N-803 for the treatment of HIV. This and similar models can inform the design of such combination therapies for cancer and chronic infection.

## Introduction

Cytokines are the chemical messengers of the immune system, controlling cell division, apoptosis, differentiation, migration, and function [1, 2]. Cytokines can be introduced therapeutically to activate the immune response against cancer [3] and chronic infections like human immunodeficiency virus (HIV) [4, 5]. IL-15 is a cytokine that promotes proliferation and function of CD8^+^ lymphocytes [6, 7]. N-803 (ImmunityBio), formerly ALT-803, is an IL-15 superagonist that, when given to non-human primates (NHPs) infected with simian immunodeficiency virus (SIV), has expanded CD8^+^ T cells [8-10], increased CD8^+^ T cell cytotoxicity [9], reduced the number of SIV infected cells in the B-cell follicles [10], and transiently reduced plasma viral load [8]. SIV is a widely used animal model of HIV with similar, albeit accelerated, disease progression in rhesus macaques [11]. N-803 also increased CD8^+^ T cell proliferation in humans participating in cancer trials [12, 13] and HIV trials [14], and it increased cytotoxicity of human CD8^+^ T cells in vitro [15]. While N-803 appears to be a promising treatment for HIV infection, the complex immunological responses to N-803 treatment remains unclear, and comparison across studies is complicated by variability in study designs. In this work, we aim to bridge experimental studies and characterize immune responses to N-803 treatment using a computational systems biology approach.

Mathematical models are a convenient complement to experimental studies and have been used to propose and evaluate hypotheses about HIV infection and the immune response for decades (reviewed in [16, 17]). These models often take the form of ordinary differential equations built upon principles of reaction kinetics. Relevant applications of these models include investigating the role of regulatory T cells in HIV infection [18], predicting the benefit of anti-PD-L1 antibody therapy [19], and analyzing viral escape from the CD8^+^ T cell response in a xenograft model of IL-15 therapy [20]. Our recent work [21] evaluated factors affecting the efficacy of N-803 in the previously mentioned cohort with transient viral suppression [8], finding that drug-induced immune inhibition, such as immune checkpoint molecule expression and regulatory T cells, could account for the observed loss of viral suppression with continued treatment.

Here we compared two cohorts of NHPs infected with SIV and treated with N-803 [8, 9]. Cohort 1 [8] had a lower pre-treatment viral load (≈3-4 CEQ/mL plasma) that was temporally suppressed with N-803 dosing, while Cohort 2 [9] had a higher pre-treatment viral load (≈5-7 CEQ/mL plasma) that was not suppressed with N-803 dosing. The reason for the distinctly different outcomes between these two studies could be connected to the degree of pre-treatment viral control [9]. In chronic HIV, higher plasma viral load is associated with increased expression of immune checkpoint molecules on CD8^+^ T cells [22, 23] and higher regulatory T cell frequency [24, 25]. Immune checkpoint molecules and regulatory T cells together form a counter-signal that limits a prolonged immune response by inhibiting CD8^+^ T cell activation, proliferation, and cytotoxic function [26-28]. These regulatory mechanisms are an important limiting factor in cytokine monotherapy aiming to promote CD8^+^ T cell control of cancer [3], including IL-15 therapy [29-31]. It is therefore possible that elevated regulatory factors in Cohort 2, induced by the higher viral load, precluded the viral suppression due to N-803 that was observed in Cohort 1. However, it is challenging to identify causal mechanisms from these experimental data alone.

In this study, we adapted and applied our computational model to simultaneously explain both the transient viral suppression in SIV Cohort 1 [8] and the lack of viral suppression in SIV Cohort 2 [9] after N-803 treatment. To this end, the mathematical model was simultaneously calibrated to SIV viral load and CD8^+^ T cell counts in both cohorts. We demonstrate how two very different viral responses to N-803 treatment can be obtained with the same model parameters if treatment is initiated at different viral loads and corresponding regulatory inhibition of CD8^+^ T cells. This work will contribute to our understanding of how the pre-treatment viral load can impact efficacy of immunotherapy.

## Methods

### Mathematical Model

We followed the convention of using ordinary differential equation models of the dynamics of viral infection and immune response (reviewed in [16, 17]). This is a lumped model that can be considered to convolve the dynamics in the blood and lymph. In this work, we updated our previous model of SIV infection and N-803 treatment [21] by separating the CD8^+^ T cell pool according to activation and SIV-specificity. This change allows the level of CD8^+^ T cell activation and regulatory inhibition to depend on both viral load and N-803, and it provides closer comparison to SIV-specific CD8^+^ T cell cytotoxic marker expression as a validation [9]. Our differential equations (Eq. 1-12) track SIV-virions (*V*), resting (*S*_0_) and active (*S*_1_-*S*_8_) SIV-specific CD8^+^ T cells, resting (*N*_0_) and active (*N*_1_,*N*_2_) non-SIV-specific CD8^+^ T cells, injection site N-803 (*X*), bioavailable N-803 (*C*), and a phenomenological representation of immune regulatory factors (*R*_1_, *R*_2_). Time dependent modifiers for proliferation (*P*), SIV-specific CD8^+^ T cell activation (*A*_*S*_), and non-specific CD8^+^ T cell activation (*A*_*N*_) are defined by algebraic equations (Eq. 13-15). Fig 1 schematically illustrates the model, all variables are defined in Table 1, and all parameters are defined in Table 2.

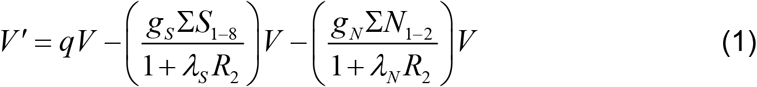

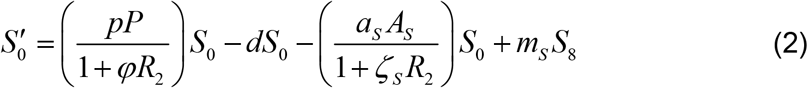

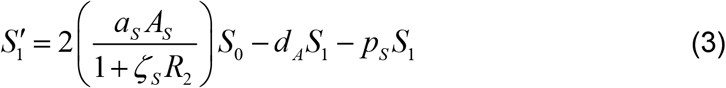

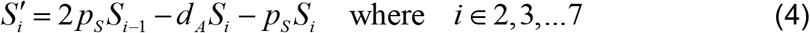

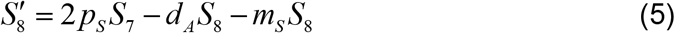

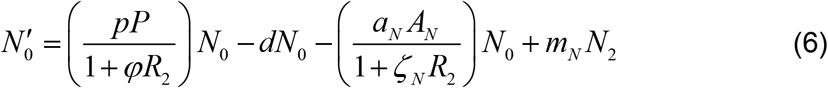

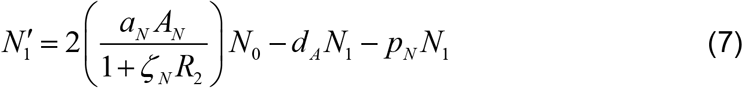

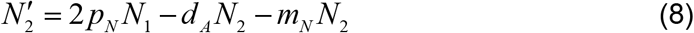

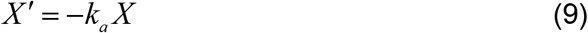

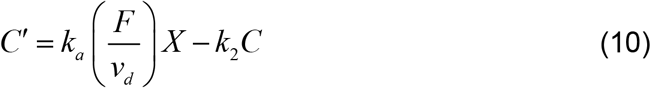

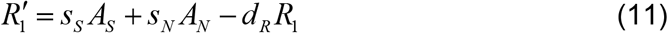

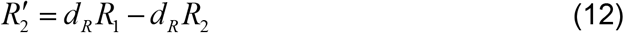

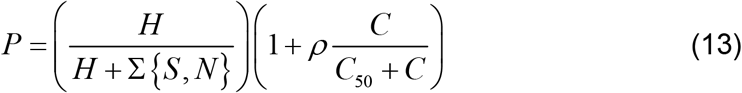

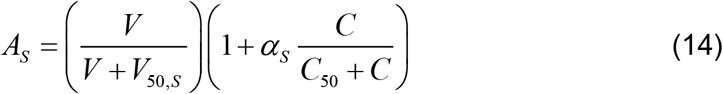

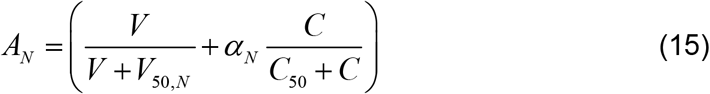

SIV virions (*V*, Eq. 1) grow exponentially (rate constant *q*) in the absence of CD8^+^ T cells. Viral growth is controlled by SIV-specific and non-specific targeting of infected cells by active CD8^+^ T cells (2^nd^ order rate constants *g*_*S*_,*g*_*N*_, respectively). Note that this simplified infection model can be obtained from a model with healthy cells and infected cells by assuming constant healthy cells and a quasi-steady-state for virions [21, 45, 46]. We did not separately model latently infected cells. The frequency of latently infected cells in chronic HIV infection is small [47], and our Cohort 1 saw only brief periods of viral control [8].

**Table 1.** Independent variables and initial conditions.

| Variable <sup>a</sup> | Symbol <sup>b</sup> | Initial Value <sup>b</sup> | Units | Ref. <sup>c</sup> |
| --- | --- | --- | --- | --- |
| SIV virions in plasma | $V$ | 3.81 (Cohort #1) | log(CEQ/mL) | [8] |
|  |  | 5.68 (Cohort #2) | log(CEQ/mL) | [9] |
| Resting SIV-specific CD8<sup>+</sup> T cells | $S_0$ | *Calculated* | #/μL | |
| Active SIV-specific CD8<sup>+</sup> T cells | $S_1$-$S_8$ | *Calculated* | #/μL | |
| Resting non-SIV-specific CD8<sup>+</sup> T cells | $N_0$ | *Calculated* | #/μL | |
| Active non-SIV-specific CD8<sup>+</sup> T cells | $N_1, N_2$ | *Calculated* | #/μL | |
| Absorption site N-803 | $X$ | 880 | pmol/kg | [32] |
| Bioavailable N-803 | $C$ | 0 | pM | |
| Regulation | $R_1, R_2$ | 1 (Cohort #1) | | |
|  |  | *Calculated* (#2) |  |  |
| CD8<sup>+</sup> T cells in peripheral blood | $\Sigma S + \Sigma N$ | 578 (Cohort #1) | #/μL | [8] |
|  |  | 435 (Cohort #2) | #/μL | [9] |
| ST8 frequency $\Sigma S / (\Sigma S + \Sigma N)$ | $f_s$ | (0.03, 0.3) (#1) | | [33-35] |
| Normalized ST8 frequency ($f_s$) | $\xi_s$ | (0.01, 1) (#2) | | |

Initial conditions are different for each cohort, with the exception of N-803 initial conditions.
<sup>a</sup> ST8 refers to SIV-Specific CD8<sup>+</sup> T cells.
<sup>b</sup> Pairs of values indicate the sampled range. Some initial conditions are calculated by assuming a steady-state prior to treatment. See S1 Appendix for details on this calculation. Symbols not
shown in Eq. 1-15 ( $f_s$ , $\xi_s$ ) are used in this initial condition calculation and defined in S1 Appendix.
<sup>c</sup> See S1 Appendix for a discussion of how parameters were obtained from given references.

**Table 2.**
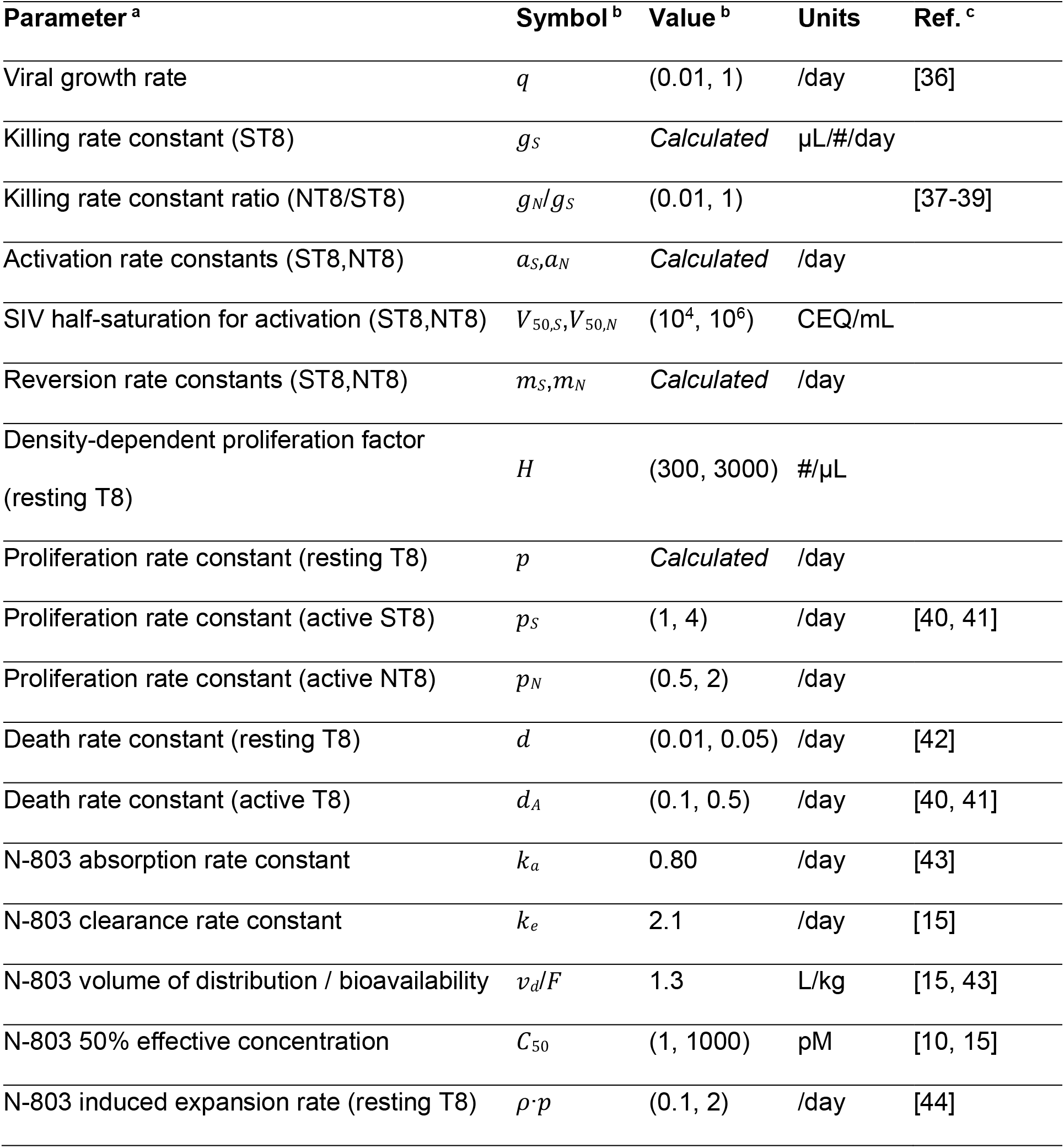
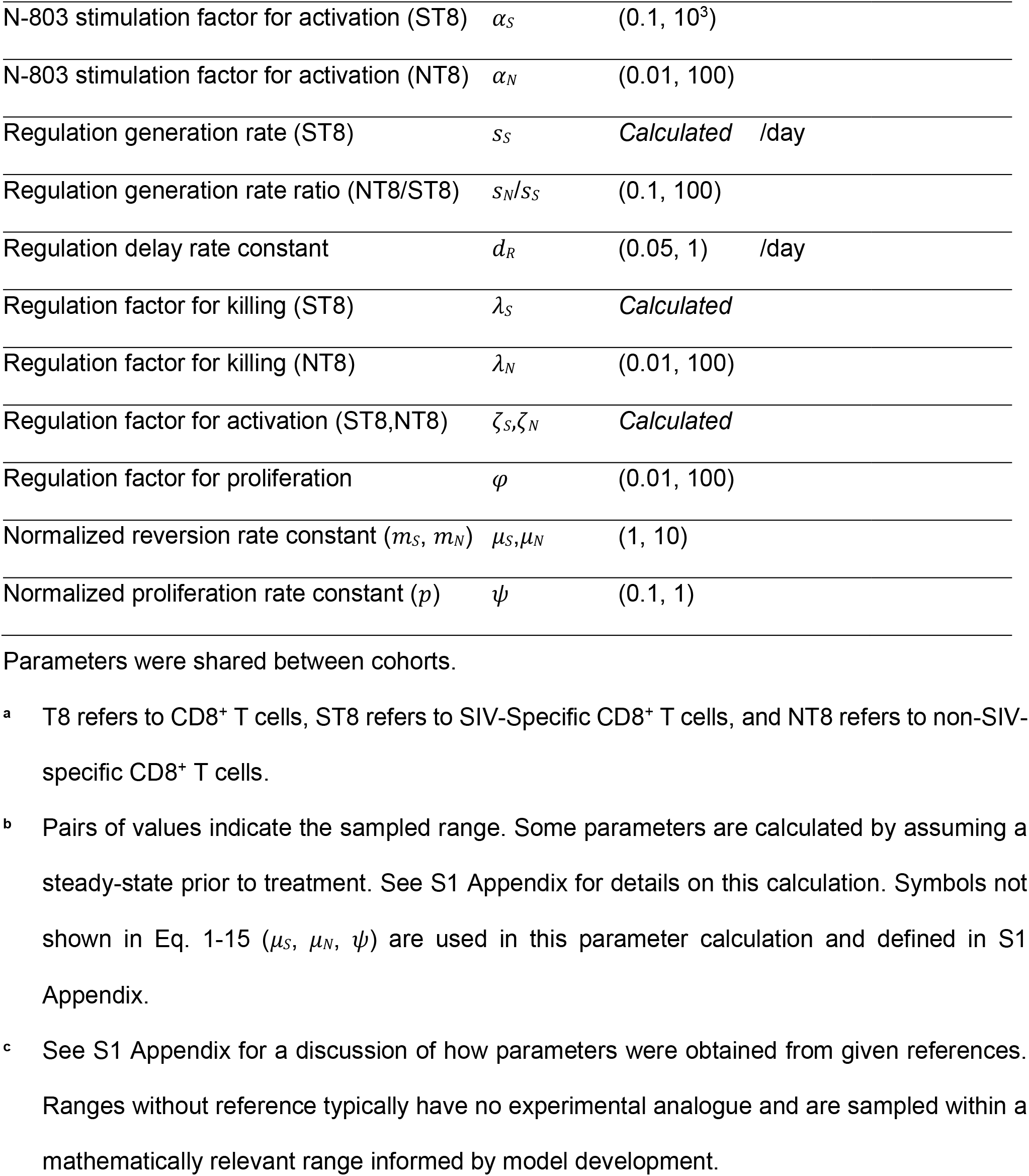
Parameter definitions and values.

**Fig 1.**
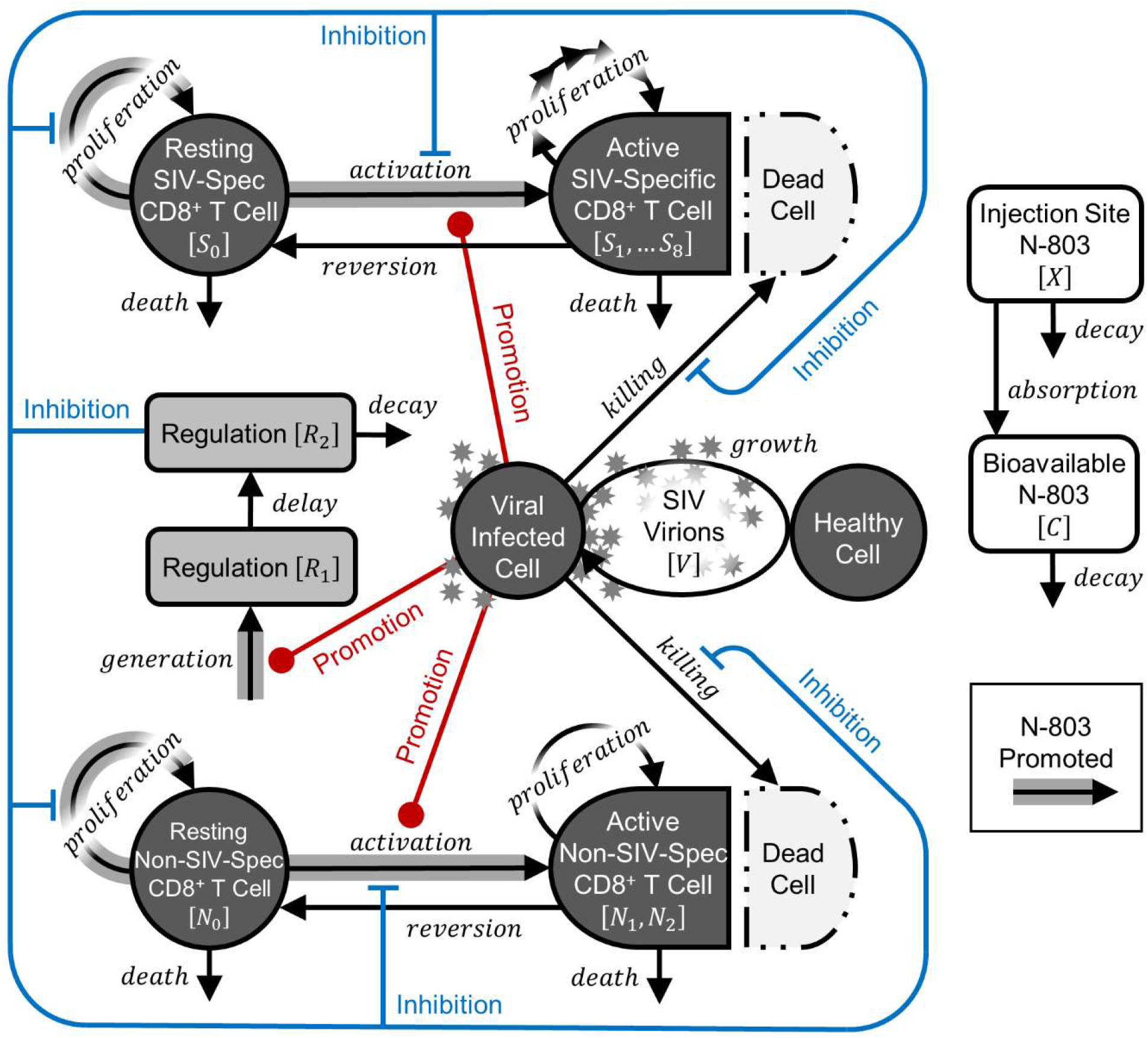
Mathematical model of SIV infection and N-803 treatment. The ordinary differential equation model was adapted from previous work [21]. SIV-virions (*V*, Eq. 1) grow exponentially and activate resting SIV-specific (*S*_0_, Eq. 2) and non-SIV-specific (*N*_0_, Eq. 6) CD8^+^ T cells. Activated SIV-specific (*S*_1_*-S*_8_, Eq. 3-5) and non-SIV-specific (*N*_1_,*N*_2_, Eq. 7,8) CD8^+^ T cells proliferate, suppress the infection, and revert to a resting state. (The chain of arrows indicates clonal expansion of SIV-specific cells.) Virus also promotes a phenomenological regulatory response (*R*_1_, *R*_2_, Eq. 11,12) that represents inhibition of CD8^+^ T cell proliferation, activation, and function by immunosuppressive cells and immune checkpoint molecules. Injection site N-803 (*X*, Eq. 9) is absorbed and becomes bioavailable N-803 (*C*), where it promotes CD8^+^ T cell proliferation and activation, as well as regulation.

Resting CD8^+^ T cells (*S*_0_,*N*_0_, Eq. 2,6) proliferate and die with a shared rate constants (*p,d* respectively), reflecting maintenance of this population by self-renewal [48, 49]. Density-dependence (*H* term, Eq. 13) is included for stability and reflects competition over space and cytokines [50]. CD8^+^ T cells are activated and divide (rate constants *a*_*S*_,*a*_*N*_ Eq. 2,3,6,7) based on saturating functions of viral load (*V*_50,*S*_,*V*_50,*N*_ terms, Eq. 14,15). Active SIV-specific CD8^+^ T cells (*S*_1_*-S*_8_, Eq. 3-5, representing 8 generations of cells) undergo expansion, which is modeled as a chain of 7 additional divisions (rate constant, *p*_*S*_), followed by reversion to a resting state (*m*_*S*_). Active cells also die faster than resting cells (*d*_*A*_). This framework is adapted from models of CD8^+^ T cell clonal expansion [40, 41]. There is also substantial non-specific activation of CD8^+^ T cells in HIV infection [51-53], and CD8^+^ T cells are capable of non-specific cytotoxicity [37]. Here, active non-SIV-specific CD8^+^ T cells (*N*_1_,*N*_2_, Eq. 7,8) are modeled similarly to SIV-specific cells, but do not undergo large, programmed expansion.

Following a standard pharmacokinetic model for subcutaneous dosing [54], N-803 (*X*, Eq. 9) is absorbed from the site of injection (rate constant *k*_*a*_). A fraction (*F*) that is absorbed becomes a bioavailable concentration (*C*, Eq. 10) over a volume of distribution (*v*_*d*_), before being eliminated (rate constant *k*_*e*_). In our pharmacodynamic model, N-803 increases CD8^+^ T cell proliferation [8, 10] and cytotoxic function [9, 15] based on multiplicative functions (Eq. 13-15) that reach half-saturation at a single concentration (*C*_50_). Note that drug and viral promotion of non-specific activation are additive with each other (Eq. 15), reflecting how IL-15 promotes non-specific activation [52]. In contrast, programmed proliferation of SIV-specific CD8^+^ T cells requires the virus (Eq. 14) but is accelerated by IL-15 [55].

Regulatory T cells and inhibitory molecules function together to prevent overreaction of the cytotoxic response (reviewed in [26-28]). We model the inhibitory effect phenomenologically with dimensionless ‘regulation’ variables (*R*_1_, *R*_2_, Eq. 11,12). Regulation is generated based on the SIV-specific and non-specific CD8^+^ T cell activation signals from the virus and N-803 (scaled by *s*_*S*_,*s*_*N*_) after a delay (rate constant *d*_*R*_), thus accounting for the effect of IL-15 on these regulatory pathways [8, 56-59]. Immune regulation acts by inhibiting CD8^+^ T cell proliferation (strength governed by *φ*, Eq. 2,6), activation (*ζ*_*S*_, *ζ*_*N*_, Eq. 2,3,6,7), and killing of infected cells (*λ*_*S*_, *λ*_*N*_, Eq. 1).

### Data Summary

Our mathematical model was simultaneously calibrated to two non-human primate (NHP) cohorts [8, 9]. These cohorts were each given N-803 during the chronic stage of a simian immunodeficiency virus (SIV) infection (Table 3). Plasma viral load, measured as viral RNA copy equivalents (CEQ), and peripheral blood CD8^+^ T cells from both cohorts were used to calibrate the model. We distinguish the cohorts by their viral load at the time of treatment, though there are other noteworthy differences. The first cohort [8], consisted of four rhesus macaques, infected with SIVmac239 and having low but detectable viral loads (≈3-4 log CEQ/mL at the time of treatment). This cohort had received vaccination against SIV prior to infection and had temporarily maintained undetectable SIV (<50 CEQ/mL) [60], and three of these macaques expressed an MHC allele associated with SIV control [61]. In the second cohort [9], fifteen rhesus macaques were infected with SIV*mac*239M, which is a barcoded SIV*mac*239 [62], and ten of these also received some type of prior vaccination. Of the fifteen NHPs, twelve had high viral loads (≈5-7 log CEQ/mL at the time of treatment). Three had viral loads near or below the detection limit, so as to make the viral load response to N-803 inconclusive for these subjects. Thus, only the twelve high viral load subjects were included in model calibration. We validated our model against measurements of Granzyme-B, a marker of cytotoxic function, on SIV-specific CD8^+^ T cells [9]. We also compare model predictions to healthy and SIV-infected NHP cohorts given intravenously administered N-803 [10].

**Table 3.** Summary of SIV Cohorts used for calibration.

| Cohort | #1 (Low Viral Load) [8] | #2 (High Viral Load) [9] |
| --- | --- | --- |
| NHP species | Indian rhesus macaques<br>(*Macaca mulatta*) | Indian rhesus macaques<br>(*Macaca mulatta*) |
| Size | 4 | 12* |
| MHC Expression | *Mamu-B\*08+* (3); *Mamu-A\*01+*<br>(1) | *Mamu-A\*01+* |
| Vaccination | All vaccinated before infection<br>[60] | 7 of 12 vaccinated before<br>infection |
| Infecting virus | *SIVmac239* | *SIVmac239M* [62] |
| Control history | Temporary viral control<br>($<50$ CEQ/mL) [60] | No history of viral control |
| Viral load at start of<br>N-803 treatment | $\approx 3\text{--}4$ log CEQ/mL | $\approx 5\text{--}7$ log CEQ/mL |
| Time post-infection<br>of N-803 treatment | 1.5 – 3 years | 6 months |
| N-803 delivery | 0.1 mg/kg subcutaneous dose<br>Once per week for four weeks,<br>one week pause, once per week<br>for four weeks, 29 week pause,<br>once per week for four weeks | 0.1 mg/kg subcutaneous dose<br>Once every two weeks for 3<br>doses total |

\* There were 15 animals in this study [9], but only 12 are used as model calibration data. Three are excluded from calibration due to their inconclusive viral response resulting from initial viral loads near or below the detection limit.

### Parameter Estimation

Maximum likelihood estimation was used to fit model outputs to plasma viral load and CD8^+^ T cell peripheral blood count in two NHP cohorts infected with SIV and treated with N-803 [8, 9]. Initial conditions were different for each cohort (Table 1), but all other parameters were shared between cohorts (Table 2). Parameter estimation was implemented by a multi-start local-search algorithm in MATLAB version R2018b (Mathworks). In brief, a Latin hypercube sample [63] of initial conditions and parameter values was generated. An interior-point algorithm [64] was used to minimize the negative log likelihood function starting from each sample. A selection of best results from the multi-start local-search algorithm were used to initialize a parallel tempering Markov chain Monte Carlo algorithm [65] to generate a Bayesian posterior distribution of parameter values. Some initial conditions and parameter values were not sampled but, instead, calculated for each sample based on the assumption that both cohorts were at separate steady-states at the onset of treatment. Details on these parameter calculations and the above calibration methods are in the S1 Appendix.

## Results

### Pre-treatment state of immune system can determine outcome of immune therapy

We simultaneously fitted our model to two SIV cohorts given N-803 (Fig 2), with Cohort 1 having a low viral load at the start of treatment (≈3-4 log CEQ/mL) and Cohort 2 having a high viral load (≈5-7 log CEQ/mL). Both of these cohorts are from previous studies and are summarized in Table 3. Fig 2A compares the simulated plasma viremia (fold change from 3.8 log CEQ/mL initial condition) to the NHP data for Cohort 1 [8] (fold change from pre-treatment mean for each subject, ≈3-4 log CEQ/mL). Following treatment initiation, the simulated viremia dropped 1.4-1.8 log within the first two weeks, compared to at least 1.3-2.1 log in NHP Cohort 1 (dropping below the detection limit of 100 CEQ/mL). The simulated viral load then rebounded to 0.5-0.8 log below the pre-treatment set point, compared to 0.5-1.3 in the experimental data, by the start of the second cycle (week 5). There was minimal response to the second cycle of doses in both the simulations and experimental data. In response to the third treatment cycle (week 37), the simulated viremia dropped by 0.5-1.2 log, with the data dropping by 0.6-1.3 log. Thus, there was a partial recovery in the efficacy of viral suppression after the long treatment break.

**Fig 2.**
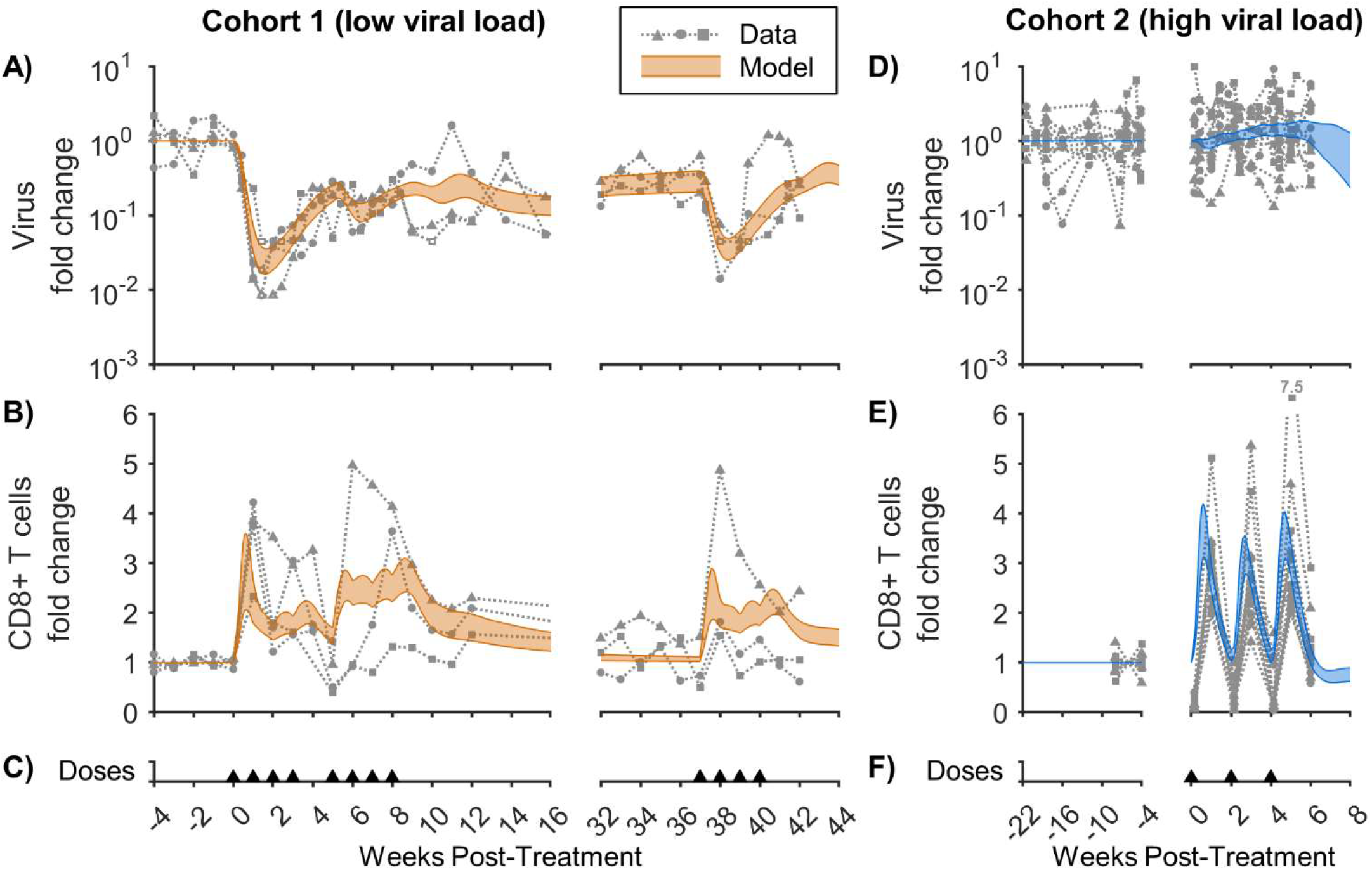
Model calibration to N-803-treated SIV cohorts with low and high viral loads. The model was calibrated to (A,D) log fold change in virus in the plasma and (B,E) fold change in CD8^+^ T cells in the peripheral blood from two different Simian Immunodeficiency Virus (SIV) cohorts. Cohort 1 (A,B) had a low viral load (≈3-4 log CEQ/mL) at the start of treatment [8], while Cohort 2 (D,E) had a high viral load (≈5-7 log CEQ/mL) [9]. The shaded region corresponds to the Bayesian 95% credible interval of the mathematical model. Open data symbols were at the lower limit of detection for the viral assay (100 CEQ/mL) and are omitted from parameter estimation. Panels (C,F) show timing of 0.1 mg/kg subcutaneous doses of N-803. Corresponding parameter distributions are shown in Fig S1 and S2 (within S1 Appendix).

Fig 2D compares the fold change of simulated plasma viremia (from 5.7 log CEQ/mL initial condition) to the fold change for Cohort 2 [9] (from pre-treatment mean for each subject, ≈5-7 log CEQ/mL). In contrast to Cohort 1, viremia did not consistently decrease in response to treatment in Cohort 2. Note that, in Cohort 1, viral load dropped precipitously (in both simulation and data) before the second dose is given, so the dosing regimens alone do not account for the differences in viral dynamics.

In contrast to viral load, CD8^+^ T cells increased in both cohorts in response to the first dose (Fig 2B,2E). CD8^+^ T cells expanded ≈2-4-fold in the simulations compared to ≈2-5-fold in the experimental data across both cohorts. In Cohort 1 (Fig 2B), the simulated CD8^+^ T cells contracted after the initial expansion, increasing again after the 1 week break in doses. This is qualitatively consistent with the data, though there was some variability in the data between individuals. In contrast, Cohort 2 (Fig 2E), with doses at 2-week intervals, showed a more consistent expansion and contraction of cells with each dose. It is worth noting that the sampling timeline is different for the two groups with Cohort 2 having data collected 1 day after each dose, and Cohort 1 having data collected immediately before each dose. A drop in peripheral blood CD8^+^ T cells in Cohort 2 was observed one day after each dose. This could be due to migration of CD8^+^ T cells from the blood to the lymph tissue [66] or mucosal tissue [67], which our model does not explicitly consider.

Taken together, our simultaneous calibration to two NHP cohorts, with shared parameters, supports the theory that pre-treatment viral load can account for starkly different viremia dynamics in response to immune therapy, despite similar CD8^+^ T cell expansion. To further support this finding, we calibrated our model to each cohort separately, to evaluate if allowing parameters to be different between the two cohorts we can achieve better model fits. Indeed, the quality of model fit did not significantly improve when fitting parameters to each cohort separately (Fig S3 within S1 Appendix).

### Model is validated by CD8^+^ T cell cytotoxicity data

To validate our model, we compared our simulated frequency of cytotoxic cells among SIV-specific CD8^+^ T cells (∑*S*_1…8_*/*∑*S*_0…8_) to experimental data that were not used in calibration (Fig 3). In the experimental data, expression of Granzyme-B among SIV-specific effector memory (EM) CD8^+^ T cells is shown [9], where Granzyme-B is a marker of cytotoxic function [68]. In our simulations, Cohort 1 (Fig 3A) started with a lower cytotoxic cell frequency, 13-46%, compared to 43-79% in Cohort 2 (Fig 3C). Pre-treatment cytotoxic frequency was similarly distinguished between the experimental cohorts, which show a frequency of 16-23% in Cohort 1 compared to 22-75% in Cohort 2. By the third day after N-803 administration, the simulated cytotoxic frequency was high for both cohorts: 78-95% in Cohort 1 and 95-99% in Cohort 2. This is compared to the experimental data showing 46-78% for Cohort 1 and 55-94% for Cohort 2. At day 7, model predicted cytotoxic frequency was similar to day 3 for both cohorts, while measured Granzyme-B frequency lowered to 45-48% in Cohort 1 and 22-79% for Cohort 2. Simulation and data were still in qualitative agreement in that SIV-specific cytotoxicity started lower in Cohort 1 than in Cohort 2 but N-803 moved the cohorts closer together. Our model was further validated by comparison to an independent dataset measuring viral load and CD8^+^ T cells after intravenous dosing in SIV-infected and SIV-naïve NHPs [10] (Fig S4 in S1 Appendix). Thus, the model was supported with both supplementary data from the cohorts of interest and from other NHP cohorts.

**Fig 3.**
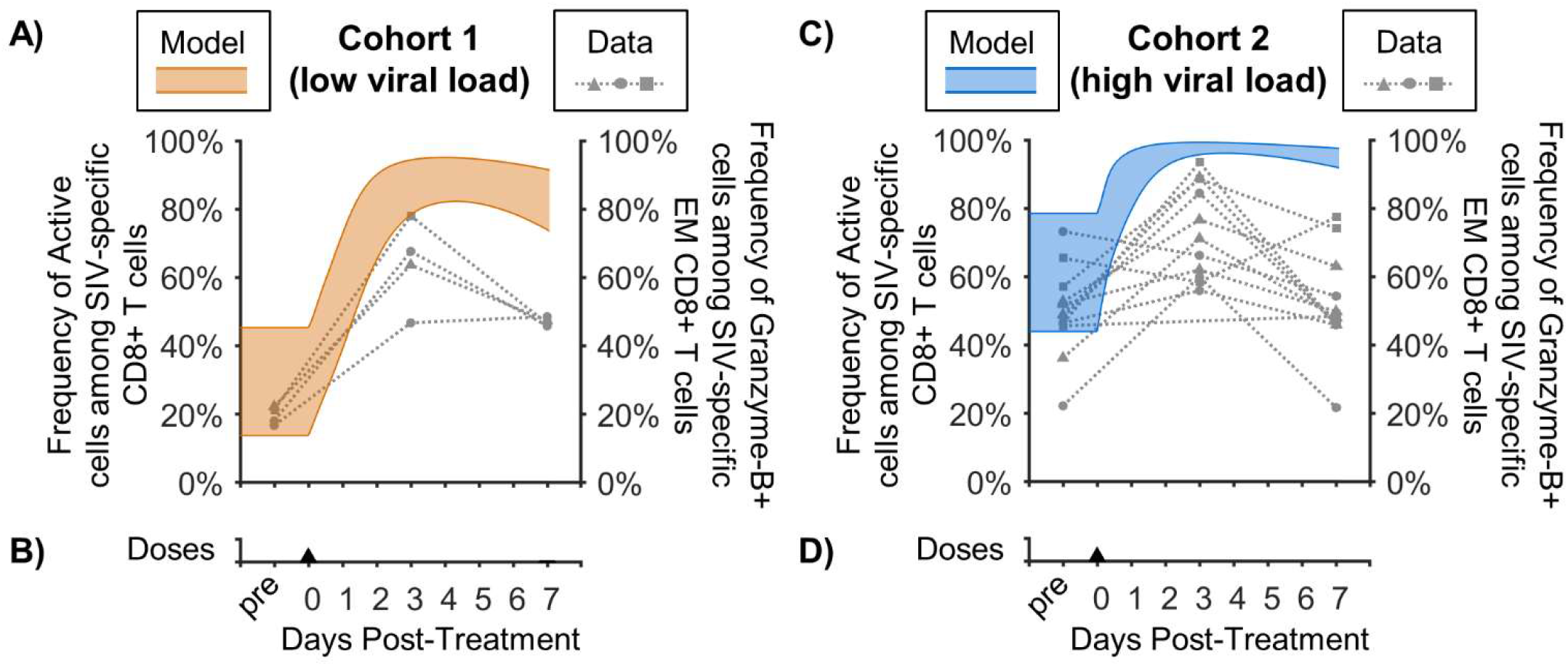
Validation of model predicted cytotoxic frequency among SIV-specific CD8^+^ cells. Shown are model predictions for the frequency of cytotoxic cells among SIV-specific CD8^+^ T cells (Bayesian 95% credible interval) for Cohort 1 (A) and Cohort 2 (C). In the model, this is the ratio of active SIV-specific CD8+ T cells to total SIV-specific CD8+ T cells (∑*S*_1…8_*/*∑*S*_0…8_). The model is compared to the frequency of Granzyme-B^+^ cells among SIV-specific effector memory (EM) CD8^+^ T cells [9] taken before treatment and at days 3 and 7 after the first dose of N-803. These cells were specific to either Gag_181-189_CM9 or Nef_137-146_RL10 epitopes, depending on primate MHC expression. Panels (B,D) show timing of 0.1 mg/kg subcutaneous doses of N-803.

### Regulatory inhibition of CD8^+^ T cell cytotoxicity precludes suppression in the high viral load cohort

Since our calibrated and validated simulations indicated that pre-treatment plasma viral load can affect treatment response, we next took advantage of our computational approach and quantitatively uncoupled the contribution of individual mechanisms to the simulated viral and immune dynamics. Fig 4 shows a selection of terms from Eq. 1-15 that are most relevant to explaining the different dynamics observed in the two cohorts. Since we were most concerned with comparing the cohorts, all values in Fig 4 are normalized to the corresponding Cohort 1 pre-treatment value. Thus, we focused on the relative change of important mechanisms and viral variables with respect to the Cohort 1 pre-treatment baseline.

**Fig 4.**
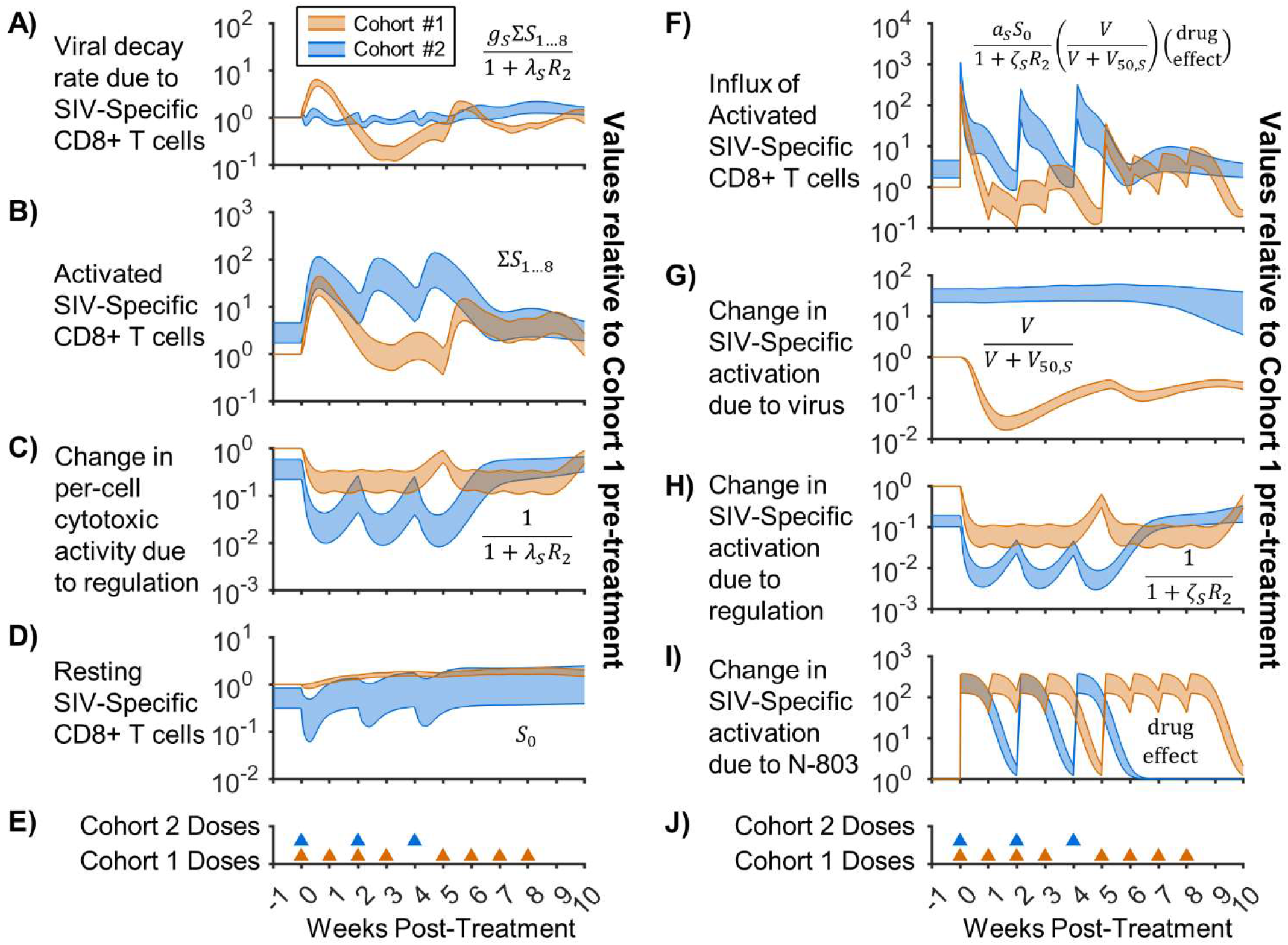
Factors contributing to viral suppression and CD8^+^ T cell activation. Shown is a breakdown of important model terms governing viral suppression and CD8^+^ T cell activation for Cohort 1 (low viral load, orange) and Cohort 2 (high viral load, blue). Values for each fitted parameter set were normalized to the Cohort 1 pre-treatment baseline for that parameter set, with Bayesian 95% credible intervals of the normalized values shown. Equivalent terms from Eq. 1-15 are shown within each axis. Panel A shows the decay rate applied to the virus by SIV-specific CD8^+^ T cells (Eq. 1), with B & C separating this term into the active SIV-specific CD8^+^ T cell count and the change in cytotoxicity due to regulatory inhibition. Panel F shows the influx of SIV-specific CD8^+^ T cells from the resting compartment (Eq. 2,3), with panel D showing the resting SIV-specific CD8^+^ T cells and panels G-I showing the contributions of antigenic stimulation, regulatory inhibition, and N-803 promotion to the activation rate. Regulation in panels C,H represents the effects of immunosuppressive cells and immune checkpoint molecules (Fig 1, Eq. 11,12). Panels E,J show the timing of 0.1 mg/kg subcutaneous doses of N-803 for each cohort.

We first evaluated the impact of SIV-specific CD8^+^ T cells on the viral load. Fig 4A shows the decay rate of the viremia due to SIV-specific CD8^+^ T cells in the model, which would be analogous to the percentage of infected cells killed per day. Figs 4B and 4C separate this rate into the contribution of active SIV-specific CD8^+^ T cell count (4B) and pre-cell reduction in killing due to immune inhibition (4C). In Cohort 1, there was a sharp increase in killing when the active SIV-specific CD8^+^ cell population increases following the first dose (Fig 4B). Although Cohort 2 has similar or greater levels of cytotoxic cells compared to Cohort 1 following dosing (Fig 4B), there was no corresponding increase in CD8^+^ T cell-dependent viral decay rate (Fig 4A). Granzyme-B must be released from the CD8^+^ T cell to kill a target cell [68], and this release can be prevented by immune checkpoint molecules [69]. We modeled this by applying inhibitory regulation to the cytotoxic function of CD8^+^ T cells (Fig 4C). The effect of inhibition was greater for Cohort 2 both before and during N-803 treatment. Since all model parameters (e.g. *λ*_*S*_) were shared between cohorts, this difference was due to higher values of the immune regulation variables (*R*_1_, *R*_2_, Eq. 11,12). Also, the difference in initial regulation between Cohorts 1 & 2 was not set arbitrarily but was rather a function of parameter values and viral load (see Table 1 and S1 Appendix for details), thus representing increased immune checkpoint molecule expression with viral load [22, 23]. In other words, the model predicts increases in cytotoxic CD8^+^ T cells in both cohorts, but the cells in Cohort 2 were rendered less effective due to inhibitory factors.

Fig 4F shows the model predicted daily influx of active SIV-specific CD8^+^ T cells from the resting state. Mathematically, this influx is a function of available antigen (viral load) (Fig 4G), regulatory inhibition (Fig 4H), N-803 promotion (Fig 4I), and the resting cells available for activation (Fig 4D). There were overlapping peaks in activation for both cohorts with the first N-803 dose, with Cohort 2 having higher activation pre-treatment. Further activation in Cohort 2 was limited by the reduced pool of resting cells (Fig 4D). Thus, for Cohort 2, there was a smaller relative benefit from N-803 due to the higher pre-treatment activation. For Cohort 1, the influx of activated cells quickly dropped with continued treatment (Fig 4F), despite strong promotion due to N-803 (Fig 4I). In the model, two factors compromised activation for Cohort 1. First was the reduction of viral load and corresponding loss of antigen (Fig 4G). Second was the increase in regulatory inhibition of activation (Fig 4H), which models potential increases in increases in regulatory T cell frequency and inhibitory molecule expression following N-803 [8]. In the model, regulatory inhibition is generated in response to increased activation, which included the effect of N-803 dosing (Eq. 11). In Cohort 1, the two 2-week break after the fourth dose allowed a partial reset of the regulatory counter-reaction, improving the activation in response to the 5^th^ dose (Fig 4F). The same is shown for the 2-week doses for Cohort 2. This theoretical benefit of 2-week dosing was demonstrated in our previous model [21].

Collectively, these results point to immune regulatory mechanisms (regulatory T cells, checkpoint molecules, etc.) as a reasonable model to explain both the lack of viral suppression following N-803 in high viral load settings (Cohort 2) and the rebound after N-803-induced viral suppression in low viral load settings (Cohort 1).

## Discussion

Here, we adapted and expanded our previous mechanistic mathematical model [21] to replicate two distinct responses to an IL-15 agonist (N-803) observed in two SIV-infected NHP cohorts. Our simulations of the two cohorts assumed the same model parameter values for both cohorts but allowed different pre-treatment steady-states. These states being the initial values of model variables representing viral load, CD8^+^ T cells, and a phenomenological amalgamation of immune inhibitory factors such as regulatory T cells and immune checkpoint molecules. The model was able to capture the CD8^+^ T cell response to subcutaneous N-803 administration in both cohorts (Fig 2B,E), the transient SIV suppression in Cohort 1 (Fig 2A), and the lack of SIV suppression in Cohort 2 (Fig 2C). The model was qualitatively validated against the experimentally measured frequency of cytotoxic cells among SIV-specific CD8^+^ T cells in both cohorts (Fig 3) and against CD8^+^ T cells and SIV viral load from independent NHP cohorts (Fig S4). This illustrates that applying an activator like IL-15 during a state of high immune activation yields a diminished return. In Cohort 2, regulatory inhibition of CD8^+^ T cell killing (Fig 4C) limited viral suppression by SIV-specific cells (Fig 4A) despite increases in cytotoxic cell counts (Fig 4B). In Cohort 1, loss of antigen stimulation (Fig 4G) combined with regulatory inhibition (Fig 4H), lowered CD8^+^ T cell activation with repeated N-803 doses (Fig 4F), allowing viral rebound.

While this work demonstrates that the divergent N-803 responses of the two cohorts can be replicated by a mathematical model where the cohorts differed by the initial state alone, there were also additional biological differences between the cohorts (Table 3). Three of the four NHPs in Cohort 1 expressed the Mamu-B*08 MHC allele [8], which has been associated with better SIV control [61]. Most notably, Cohort 1 temporarily controlled SIV as part of a previous study [60]. The biological mechanisms behind spontaneous HIV control are still a subject of much research [70, 71]. Inherent differences between the cohorts could be represented mathematically by different model parameter values, which could allow for theoretical assessment of potential mechanisms of HIV control. Here we illustrated that these inherent biological differences are not mathematically necessary in order to recreate the observed viremia dynamics.

Both this current work and our previous work [21] theoretically demonstrate how immune regulatory mechanisms could be responsible for the lack viral suppression by N-803. Since N-803 seems to be effectively expanding cytolytic SIV-specific cells in both cohorts (Fig 4B), direct reduction of the per-cell killing rate of cytotoxic cells (Fig 4C) is necessary to preclude viral suppression in the mathematical model. Biologically, this reduction in cytotoxic activity could be caused by shielding of infected cells via PD-L1 on these cells, signaling through PD-1 on cytotoxic cells and stopping the cytolytic response [69]. A combination of N-803 with two anti-PD-L1 domains (N-809) was able to promote the cytotoxic function of CD8^+^ T cells and natural killer cells as well as bind to tumor cell PD-L1, synergistically increasing tumor cell lysis in vitro and in murine models [30, 31]. In addition, N-809 reduced the frequency of immunosuppressive regulatory T cells (defined as CD4^+^ FoxP3^+^ T cells) in the tumor environment [31]. In a mouse model of chronic lymphocytic choriomeningitis virus, depletion of regulatory T cells (also CD4^+^ FoxP3^+^) expanded virus specific CD8^+^ T cells but did not reduce viral load, possibly due to a simultaneous increase in PD-L1 on target cells [72]. When regulatory T cell depletion and PD-1 blockade were simultaneously applied in the mouse model, viral load was reduced. It is conceivable that combining N-803 with regulatory T cell depletion may be redundant, as both would increase virus-specific CD8^+^ T cell numbers while reducing their killing efficacy by increasing PD-1/PD-L1 expression.

Similar combinations of immune activators, like N-803, and immune checkpoint inhibitors are being included in multi-drug therapy of cancer. In cancer, chronic inflammation promotes expression of immune checkpoint molecules and their ligands [73, 74], which limit the effectiveness of IL-15 therapy [30, 31, 75]. Inflammation also recruits immunosuppressive cells, such as myeloid-derived suppressor cells (MDSCs) [76], which, in turn, attenuate the effectiveness checkpoint blockade therapy [77, 78]. In light of these many factors, increasingly complex therapies are being proposed. A murine tumor model demonstrated the effectiveness of a six-drug combination therapy that included N-803 and an anti-PD-L1 antibody [79]. A recent case study saw complete remission of Merkel cell carcinoma following a combination of N-803, PD-1 blockade, and a chemotherapeutic [80]. Both of these regimens included taxanes, which are anti-mitotic chemotherapy drugs that also promote antigen presentation in cancer cells to enhance targeting by CD8^+^ T cells [81]. Some chemotherapeutics can also target MDSCs [76, 82], and combining N-803 with sunitinib to suppress MDSCs synergistically inhibited melanoma growth and promoted survival in a murine model [83]. Treatment of certain types of tumors may require these multifaceted approaches [79], and mechanistic mathematical models, such as the one developed here, could be developed to inform the integration of these approaches and design regimens that may have improved efficacy.

In HIV, the approach outlined above would be somewhat analogous to combining immune therapy with traditional antiretroviral therapy (ART). ART can reduce PD-1 expression in CD8^+^ T cells [84, 85], and reduce the frequency of CD4^+^ FoxP3^+^ regulatory T cells [86, 87]. Comparison of CD8^+^ T cell proliferation and cytotoxicity markers between these two cohorts supported the assertion that N-803 is best used in the context of patients that control viral loads [8]. Though a recent clinical trial demonstrated N-803 can be safely administered to ART-suppressed HIV patients [14], combination of N-803 and ART in an animal cohort did not reduce the latent SIV reservoir or preclude viral rebound after ART interruption [88]. There is still the potential that the further addition of PD-1 blockade to ART and N-803 combination therapy could allow viral control with less frequent dosing. Such a switch from daily to weekly regimens for the treatment of osteoporosis improved patient adherence [89, 90]. There are additional mechanisms that would be relevant for a mathematical model to consider application of N-803 to ART-suppressed HIV. First is the capacity of N-803 to reverse latent HIV infections [88, 91]. Latency was excluded from this model under the assumption that reactivation of latent SIV would only be a small contribution to unsuppressed viral loads, given the small frequency of latently infected cells in chronic HIV infection [47]. Second, N-803 can induce CD8^+^ T cell infiltration into B cell follicles [10], which are sites with dendritic-cell-bound viral reservoirs and limited CD8^+^ T cell presence [92].

In summary, we used a mathematical model, informed by in vivo treatment data, to demonstrate that the response to a therapeutic cytokine will depend on the pretreatment immune state and the balance between immune activation and regulation. These results can provide further insights into the application of IL-15-based therapy in the treatment of both cancer and chronic infections.

## Acknowledgments

We thank ImmunityBio for supplying the reagent N-803.

## S1 Appendix

### Parameter Space and Fitted Distributions

The following is a discussion of the initial conditions in Table 1. Pre-treatment steady state viral load and CD8^+^ T cell numbers are calculated from experimental measurements in the two NHP cohorts. Initial SIV plasma viral loads are the means of pre-treatment data points, taken after converting to log(CEQ/ml), for each cohort (20 samples total for Cohort 1 [1], 78 samples for Cohort 2 [2]). We take SIVmac239 gag copy equivalents (CEQ) in the plasma as proportional to SIV virions. Initial CD8^+^ T cells in peripheral blood are the means of pre-treatment data points for each cohort (20 samples total for Cohort 1 [1], 24 samples for Cohort 2 [2]). The frequency of HIV-specific CD8^+^ T cells in humans with HIV is variable (≈1-20%) [3-5]. We expand this range upward to account for potential differences in SIV.

The normalized frequency of SIV-specific CD8^+^ T cells for cohort 2 (*ξ*_*S*_) is defined in the *Parameter Calculation* section. Since only the effect of immune regulation is relevant, the values of the regulation variables (*R*_1_, *R*_2_) are normalized to the pre-treatment value for Cohort 1. Thus, regulation strength parameters (*φ, λ*_*S*_, *λ*_*N*_, *ζ*_*S*_, *ζs*_*N*_) reflect the pre-treatment effect of regulation for Cohort 1. The initial pmol/kg of N-803 at the absorption site (*X*) for a 0.1 mg/kg dose of N-803 is obtained from the measured molecular weight of 114 kDa [6].

The following is a discussion of the parameter values in Table 2 that govern viral infection and the CD8^+^ T cell response. Viral growth rate (*q*) is based on estimates of HIV-infected cell death rate due to CD8^+^ T cells (reviewed in [7]), since *q* balances the effect of CD8^+^ T cells in the pre-treatment steady-state. Non-specific CD8^+^ T cell cytotoxic function is similar to natural killer cells [8], so their rate of killing (*g*_*N*_) is assumed a fraction of SIV-specific CD8^+^ T cell rate of killing (*g*_*S*_). This follows from studies where depletion of CD16^+^ natural killer cells during acute infection had a negligible effect on viral load [9], while depleting all CD8^+^ lymphocytes during chronic infection increased viral load by up to 1000 fold [10]. The proliferation rate of active SIV-specific CD8^+^ T cells (*p*_*S*_) reflects estimates used in CD8^+^ T cell clonal expansion models [11, 12]. Influenza-specific memory CD8^+^ T cell cycle in 6 hours [13] in response to antigen, which translates to a proliferation rate constant of *p*_*S*_ = 2.77/day. Proliferation for active non-SIV-specific CD8^+^ T cells (*p*_*N*_) is assumed to be slower. Active cell death rate (*d*_*A*_) is sampled based on models to CD8^+^ T cell clonal expansion [11, 12], and the resting CD8^+^ T cell death rate (*d*) is based on measured turnover in healthy monkeys [14]. Proliferation stimulation factor (*ρ*) is limited according to a maximum drug-induced expansion rate for the SIV-naive case (*p*·*p* for this model), which is assumed to be less than the CD8^+^ T cell clonal expansion rate in rhesus macaques (≈1/day) [15]. Other pharmacokinetic and pharmacodynamic parameters (*k*_*a*_, *k*_*e*_, *v*_*d*_*/F, C*_50_) are carried over from our previous work [16]. The normalized parameters (*μ*_*S*_, *μ*_*N*_, *ψ*) are defined in the *Parameter Calculation* section.

Figures S1 shows the distributions resulting from Bayesian MCMC for the parameters of Eq. 1-15. Figures S2 shows the distributions for the parameter convolutions sampled and used to calculate certain parameters.

**Figure S1.**
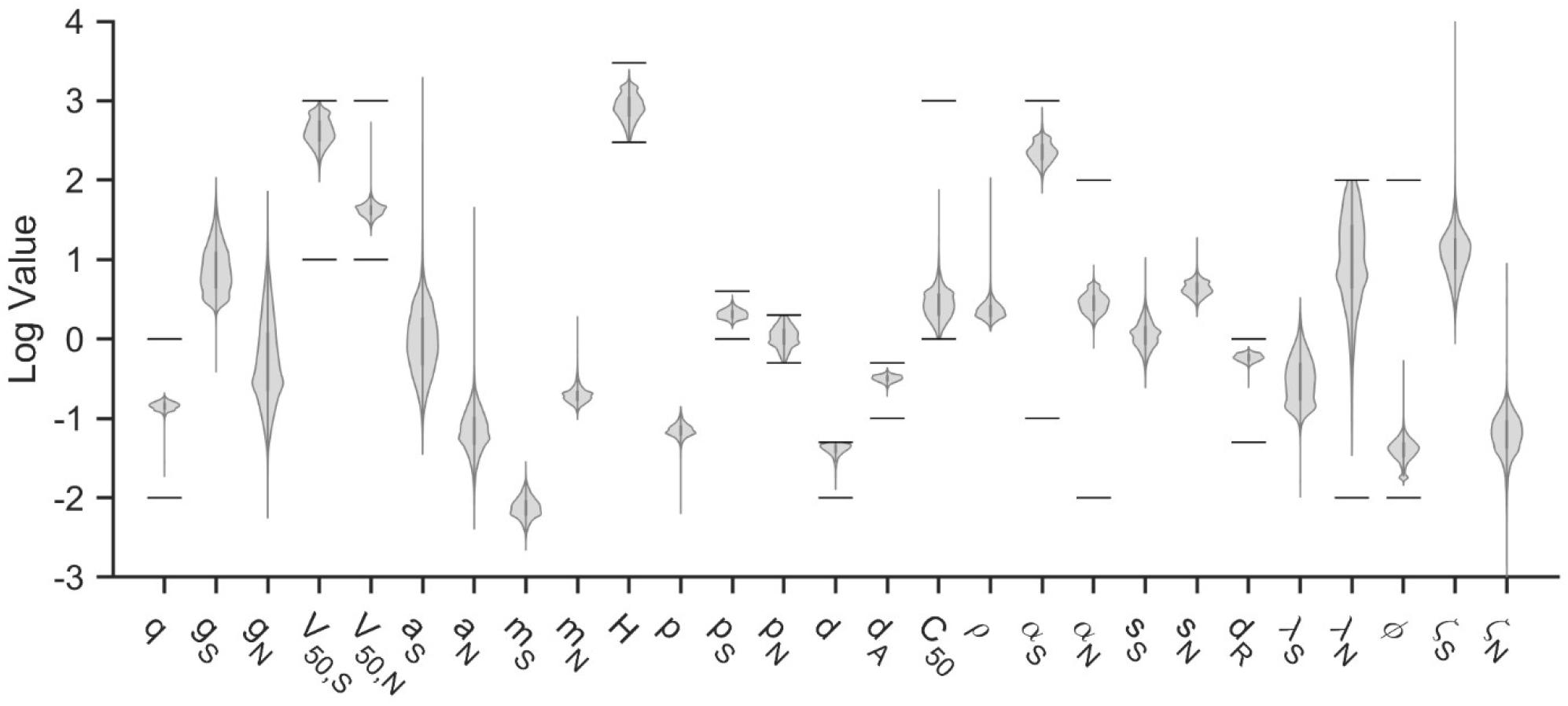
Fitted parameter distributions. Shown is the Bayesian MCMC sample of parameter values from Eq. 1-15. Solid lines indicate allowed parameter ranges, which are also given in Table 2. Parameters without ranges are calculated by assuming pre-treatment steady-state. Distributions of sampled parameter convolutions used in those calculations are shown in Fig S2. Units are as given in Table 2, with the exception of *V*_50,*S*_, *V*_50,*N*_, which are converted to CEQ/μL.

**Figure S2.**
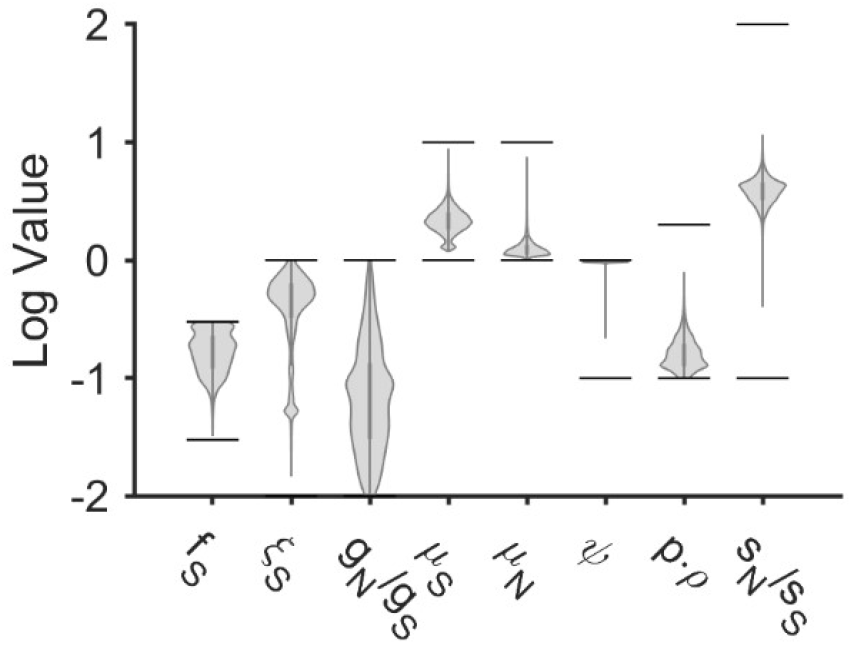
Fitted distributions for parameter convolutions. Shown is the Bayesian MCMC sample of parameter convolutions used to calculate model parameters and initial conditions in order to maintain a pre-treatment steady-state. Solid lines indicate allowed ranges, which are also given in Table 2. Units are as given in Table 2. The normalized SIV-specific cell frequency (*ξ*_*S*_), reversion rate constants (*μ*_*S*_, *μ*_*N*_), and resting proliferation rate constant *ψ*(*ψ*) are defined in Eq. S35, S24, and S21,S22, respectively.

### Parameter Calculation

Prior to treatment, the system is approximated to be at steady-state. The ODEs become equations that relate initial conditions. Note that the definitions of modifiers *P, A*_*S*_, *A*_*N*_ (Eq. 13-15) have been replaced with total initial rates *p*^*^, 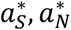 (Eq. S11-14). This is for convenience with the proceeding analysis.

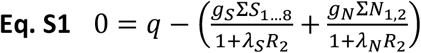

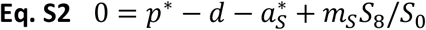

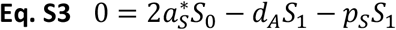

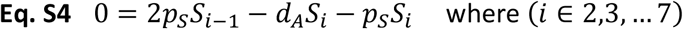

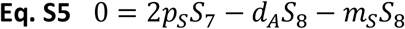

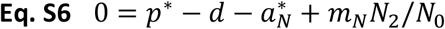

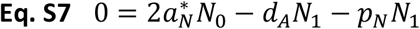

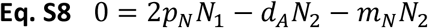

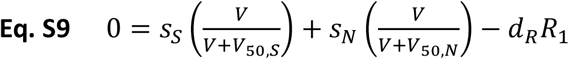

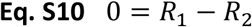

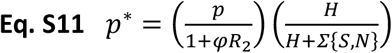

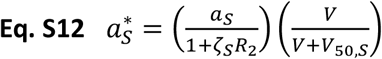

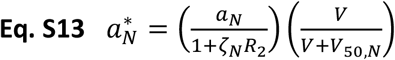

The equations below describe how unknown initial conditions and parameter values are obtained. Special care is taken to avoid negative parameter values. This effort is not foolproof, so parameter sets with negative values are filtered out of the MCMC sample (<0.3%).

#### Regulation initial values and generation parameters: [*R, s*_*S*_, *s*_*N*_]

Regulation is normalized to the low-VL group (*R*_*l*_ *= 1*). Regulation for the high-VL group (*R*_*h*_) is calculated (Eq. S14), where the low and high initial viral loads are *V*_*l*_ and *V*_*h*_ and where the ratio *s*_*N*_*/s*_*S*_ is sampled.

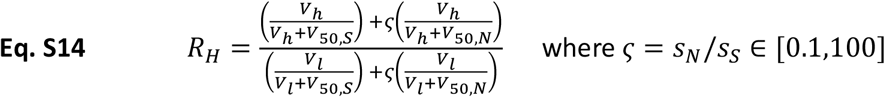

Regulation generation parameters [*s*_*S*_, *s*_*N*_] can be calculated from either group (shown below for low-VL).

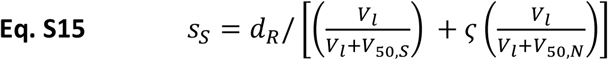

#### Reversion rate constants: [*m*_*S*_, *m*_*N*_]

A relationship between initial activation rates 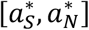 can be used to inform that values for [*m*_*S*_, *m*_*N*_]. Eq. S3-S5 relate *S*_0_ and *S*_8_, and Eq. S7-S8 relate *N*_0_ and *N*_2_. (Ratios [*U*_*S*_, *U*_*N*_] are defined because they appear frequently.)

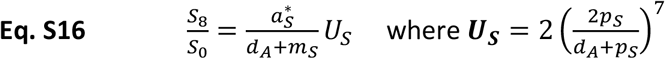

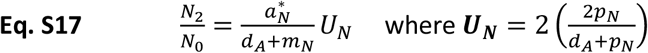

These can be combined with Eq. S2 and Eq. S6 to find the ratio between initial activation rates 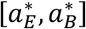.

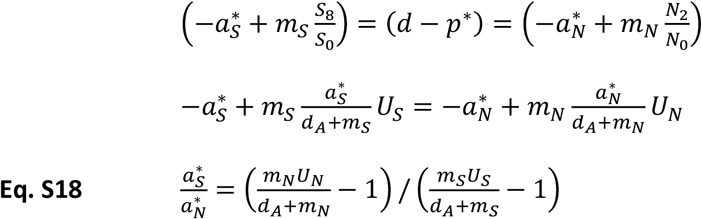

To avoid negative activation rates, both the numerator and denominator of Eq. S18 must be the same sign. One way of achieving this is to ensure that both of the following inequalities hold.

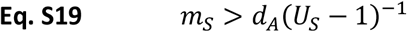

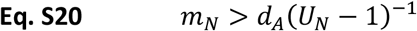

Thus, we defined **normalized reversion rate constants (*μ***_***S***_, ***μ***_***N***_), sampled as shown.

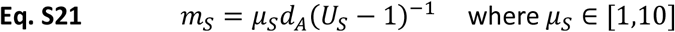

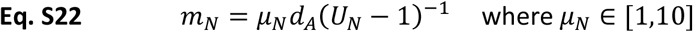

#### Resting proliferation rate constant: *p*_0_

If we consider that *p*^*^ − *d* = 0 in the uninfected (no activation or regulation) steady-state, it follows that *p*^*^ − *d* < 0 in the infected steady-state, because regulation would reduce *p*^*^.

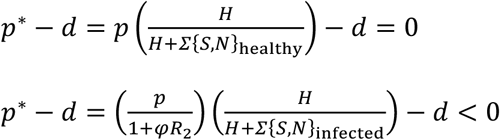

Thus, *p* must satisfy the following inequality for both cohorts.

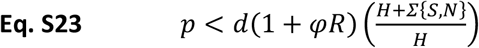

We define and sample the **normalized proliferation rate constant (*ψ*)** and use it to calculate *p* (Eq. S24), using the smaller of the values from each cohort.

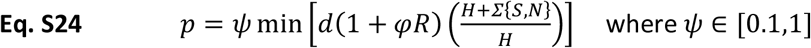

#### Activation rate constants and regulation factors: [*a*_*S,O*_, *a*_*N,O*_, *a*_*S,2*_, *a*_*N,2*_]

Once again combining Eq. S2,S6 with Eq. S16,S17, we can solve for initial activation rates 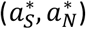.

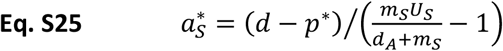

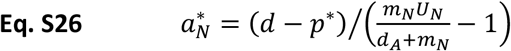

Note that, since *p*^*^ is different for each cohort (different regulation and cells), 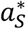 and 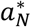 are also different. The following calculations are done for both S and N based on the definitions of 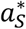 and 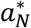 (subscripts dropped).

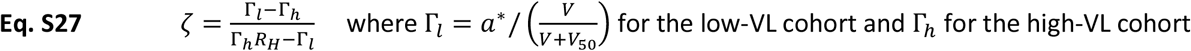

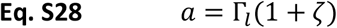

#### Initial conditions for CD8^+^ T cell subgroups: [*S*_O,…8_, *N*_O,…2_]

Eq. S3-S5 and S7-S8 can be used to relate the sum of activated *S,N* (*S*_*A*_,*N*_*A*_) to 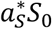 and 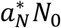 (ratios ***Z***_***S***_, ***Z***_***N***_).

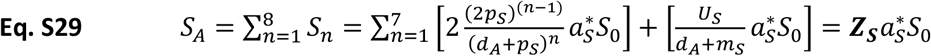

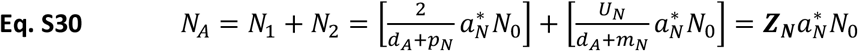

For each cohort, the initial total CD8^+^ T cells are known, and the **fraction of specific cells (*f***_***S***_) is sampled.

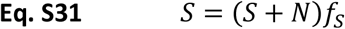

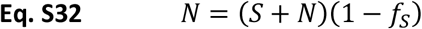

Since we now know *S*_*A*_/*S*_0_ and *N*_*A*_/*N*_0_, we can calculate *S*_0_ and *N*_0_.

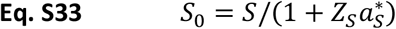

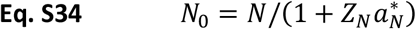

The activated subgroups then follow easily from Eq. S3-S5 and S7-S8.

#### Application of normalized specific cell fraction : *ξ*_*S*_

The scheme above does not wholly eliminate the possibility of negative parameter values. It was found that cases where the initial fraction of SIV-specific CD8^+^ T cells *f*_*E*_ was higher in cohort 1 and in cohort 2, negative parameter values would often result. Thus, we assume that *f*_*S,h*_ > *f*_*S,l*_ and used the following definition to determine *f*_*S,h*_ from sampled *f*_*S,l*_. (Note that the 0.3 comes from the maximum value for *f*_*S*_.)

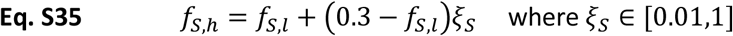

#### Killing rate constants and regulation factors: [*g*_*S*_, *g*_*N*_, *λ*_*S*_]

If the non-specific killing regulation factor (*λ*_*N*_) and the killing rate ratio *g*_*N*_/*g*_*S*_ are sampled, then *g*_*S*_ can be calculated from the following quadratic equation.

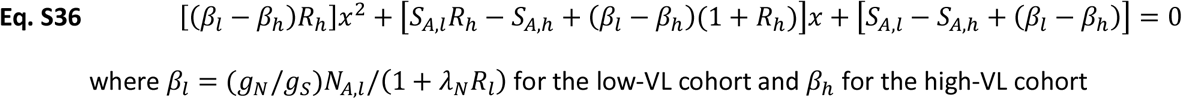

Base SIV-specific killing rate (*g*_*S*_) can then be obtained from Eq. S1.

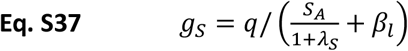

### Analytical Methods

The goal of this work is to test a hypothesis that responses to immune therapy can depend upon the state of the immune system, rather than make individualized predictions. Thus, we fit the model to collective cohort data, rather than to each individual subject, to not overinterpret individual variation and noise. There are four response variables, virus and cells from each of two cohorts. In the error model (Eq. S38), response variables (indexed by *i*) are normalized to the initial conditions for each individual NHP (indexed by *j*), which is estimated as the mean of the pre-treatment data points. Virus fold change is also log-transformed. We apply a constant error model to these transformed variables, assuming independent, identical, and normally distributed error 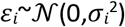 for each and neglecting error covariance between virus and CD8^+^ T cells. Note that *E* corresponds to total CD8+ T cells (∑*S*_0…8_+∑*N*_0…2_).

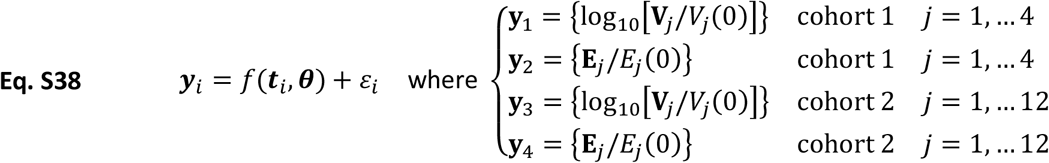

The concentrated likelihood method [17] allows the error variance a 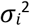 to be eliminated from the negative log likelihood function (Eq. S39), which is then a function of the sum of squared error, *S*_*i*_, and the number of data points, *n*_*i*_, for each response variable. The few viral data points at the lower limit of detection for the viral assay (100 CEQ/mL) are omitted from the likelihood function.

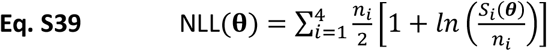

The parameter samples in Figures S1 and S2 are obtained from the ranges in Tables 1 and 2 by a Multi-Start Local Search followed by a Bayesian Markov Chain Monte Carlo (MCMC) implemented in MATLAB version R2018b (Mathworks). First, 2000 parameter samples are generated by Latin hypercube [18] (MATLAB’s ‘*lhsdesign*’), assuming loguniform distributions. Second, each sample set of parameters is optimized using an interior-point algorithm [Byrd00] (MATLAB’s ‘*fmincon*’), using logarithmic parameter values and minimizing Eq. S39. Finally, a selection of 10 parameter sets with comparable NLL and quality of fit initialize a parallel tempering MCMC algorithm [19] (*PEST*O toolbox [20]), which samples from Bayesian posterior distributions of parameter values using Eq. S39 and assuming loguniform prior distributions of parameter values. After one million iterations, the final sample is thinned by selecting very 100^th^ parameter set, yielding a 10,000-set sample. From these, a small number (<0.3%) were excluded for having negative parameter values. The remainder was used for figures and statistical analyses.

ODEs were solved using MATLAB’s stiff equation solvers, with settings dependent on the task. For parameter estimation and uncertainty quantification, we used *ode23s* with looser error tolerances (1% relative tolerance, 0.001 absolute tolerance). Tolerances were chosen by comparing model outputs over the time interval of the NHP data, with 1% relative tolerance yielding no appreciable difference from tighter tolerances. In *ode23s*, absolute tolerance is the lowest model output value for which error is considered, which we set based on a trial calibration of the model. For plotting, we used *ode1*5*s* with default error tolerances (0.1% relative tolerance, 10^−6^ absolute tolerance). In both cases, an analytical Jacobian function was provided to accelerate solutions and improve calibration outcomes.

### Single Cohort Parameter Estimation

The entire fitting process is repeated for each cohort individually to see if substantively better fits could be obtained. With only one pre-treatment steady-state to consider, fewer parameters have to be calculated. Additional parameters sampled are summarized in Table S1. SIV-Specific CD8^+^ frequency *f*_*S*_ is sampled for each cohort (per Table 1), and regulation is normalized to pre-treatment state. Instead of sampling the normalized proliferation rate constant (Eq. S24), the activated fraction (*f*_*S*_ = ∑S /∑S) for SIV-specific CD8^+^ T cells is sampled and used to calculate the equivalent fraction for non-SIV-specific CD8^+^ T cells using the scheme below.

Eq. S4,S5,S8 can also be used to relate S_*A*_ and N_*A*_ to terminally proliferated *S*_8_ and *N*_2_ (ratios ***W***_***S***_, ***W***_***N***_).

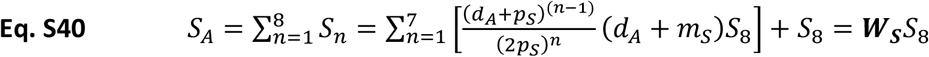

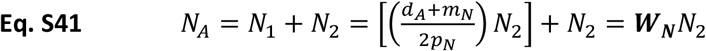

These ratios can be combined and used to write Eq. S2 and S6 in different forms (ratios ***Q***_***S***_, ***Q***_***N***_).

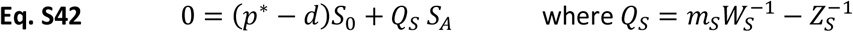

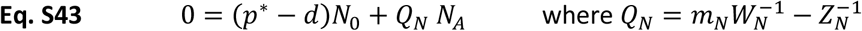

If we define and sample 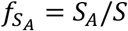, then we can use [*Q*_*S*_, *Q*_*N*_] to calculate 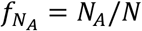.

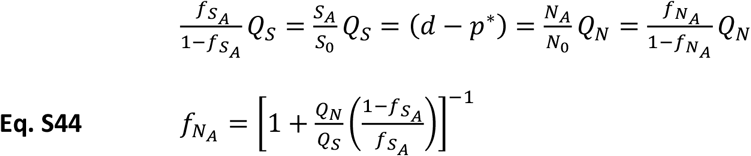

Results of calibration with and without shared cohort parameters are compared in Figure S3. Though there are small differences, allowing parameters to vary between cohorts does not improve quality of fit to the NHP data in any meaningful way.

**Table S1.** Initial Condition and Parameters for Single Cohort Fitting.

| Initial Condition or Parameter | Symbol | Value |
| --- | --- | --- |
| Active ST8 frequency $\Sigma S_A / \Sigma S$ | $f_{SA}$ | (0.009, 0.9) |
| Regulation factor for activation (ST8,NT8) | $\zeta_S, \zeta_N$ | (0.01, 100) |
| Regulation factor for killing (ST8) | $\lambda_S$ | (0.01, 100) |

**Figure S3.**
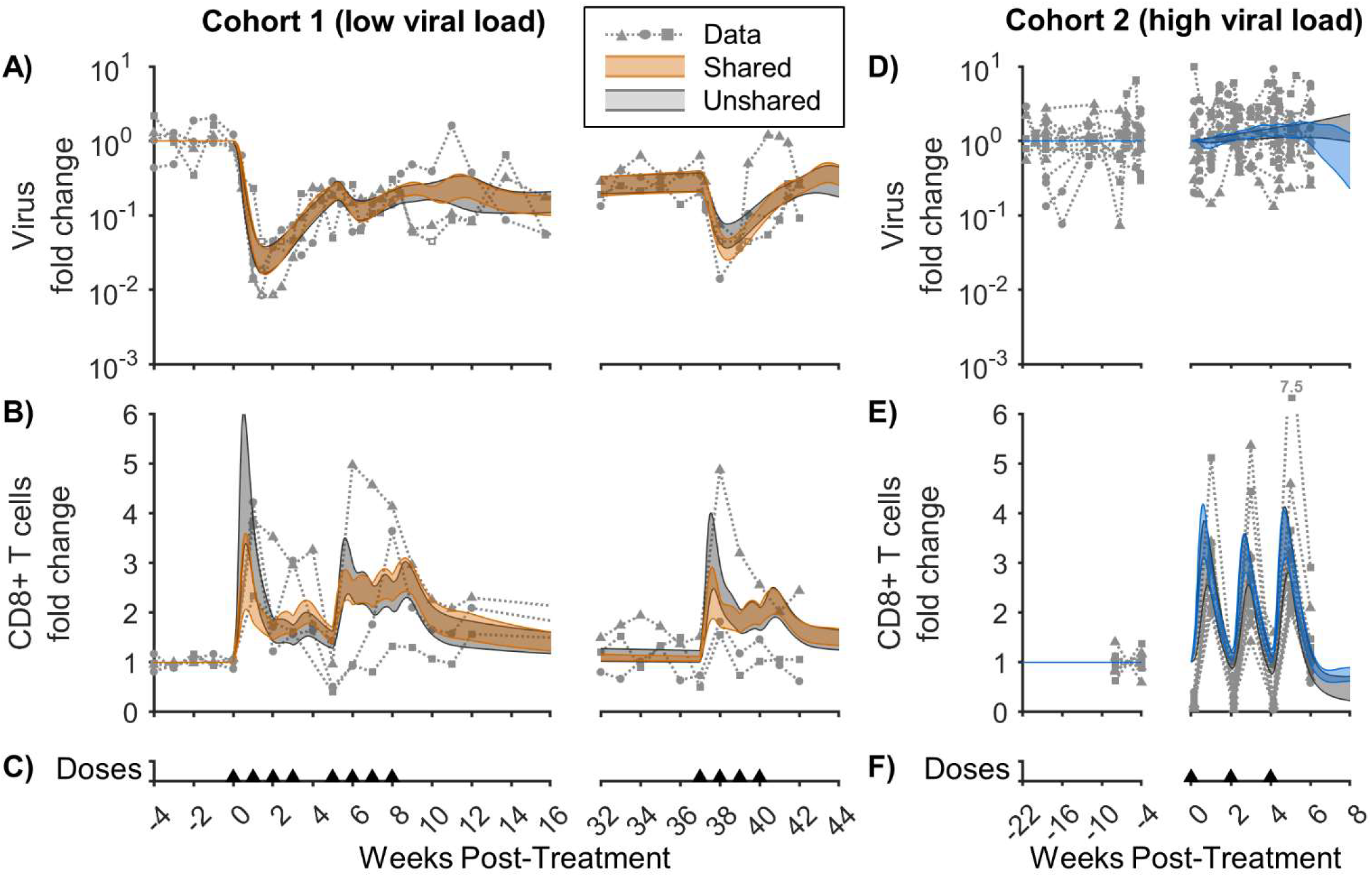
Comparison of fitting to single cohorts and fitting to both cohorts. The model was calibrated to (A,D) log fold change in virus in the plasma and (B,E) fold change in CD8^+^ T cells in the peripheral blood from two different Simian Immunodeficiency Virus (SIV) cohorts. Cohort 1 (A,B) had a low viral load (≈3-4 log CEQ/mL) at the start of treatment [1], while Cohort 2 (D,E) had a high viral load (≈5-7 log CEQ/mL) [2]. The gray shaded regions corresponds to the Bayesian 95% credible interval of the mathematical model when parameters are allowed to vary between cohorts (“Unshared”). The superimposed orange and blue shaded regions correspond to Figure 2, where the model parameters for each cohort are the same (“Shared”) but the initial conditions differ. Open data symbols were at the lower limit of detection for the viral assay (100 CEQ/mL) and are omitted from parameter estimation. Panels (C,F) show timing of 0.1 mg/kg subcutaneous doses of N-803.

### Validation Against Other Cohorts

The model was also validated against intravenously administered N-803 in SIV-naïve and SIV-infected non-human primates [21]. To model intravenous administration, doses were applied directly to the bioavailable compartment. Thus, *X*_0_/*v*_*d*_ became the initial condition for the compartment, which decayed at the same elimination rate (*k*_*e*_, Eq. S45).

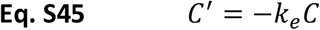

To model the SIV-naive case (Fig S4A), SIV virions (*V*, Eq. 1) and SIV-specific CD8^+^ T cells (*S*_0_-*S*_8_, Eq. 2-5) were absent. Also, active non-SIV-specific CD8^+^ T cells (*N*_1_, *N*_2_) and immune regulation (*R*_1_, *R*_2_) were initially zero but could still be induced by N-803. Thus, resting non-SIV-specific CD8^+^ T cells (*N*_0_) composed the entire CD8^+^ T cell pool prior to treatment. The pre-treatment steady-state of these cells was calculated from parameters (Eq. S46). Thus, initial conditions in the model do not directly match the initial values in the validation data, as this was necessary to have a pre-treatment steady-state using the same parameter sets calibrated to Cohort 1 [1] and Cohort 2 [2].

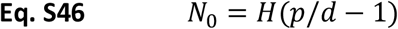

Figure S4A shows the CD8^+^ T cells in the blood of SIV-naive NHPs (n=4) after a 6 mg/kg intravenous dose of N-803 [21]. There is a brief drop in CD8^+^ T cells at day 1. This could be due to extravasation [22, 23], which that our model does not incorporate. Otherwise, both model and data show a modest expansion of CD8^+^ T cells. This can be compared to the larger expansion after an identical dose given to SIV-infected NHPs (also n=4, Fig S4D). Figure S4D also shows the results of two additional 6 mg/kg intravenous doses given at 1 and 2 weeks after the first dose, as well as a large 100 mg/kg dose at week 7. Both model and data show smaller expansions after the two subsequent 6 mg/kg doses than after the first dose (week 0). Model and data seem to diverge for the 100 mg/kg dose, where the model predicted a weaker expansion than is observed in the data. This could be due to the reduced viral load in the model (Fig S4C).

Figure S4C shows the response of viral load for one subject after the 100 mg/kg intravenous dose of N-803 [21], being the only subject with viral load data that was above the detection limit. Both model and data show an approximately 10-fold reduction in viral load following this dose, and both return to the pre-dose state after a brief period. Thus, our model reproduces several key qualitative aspects of an N-803 data set with an independent NHP cohort, different disease state (SIV-naïve vs. SIV-infected), and different route of N-803 delivery (intravenous vs. subcutaneous).

**Figure S4.**
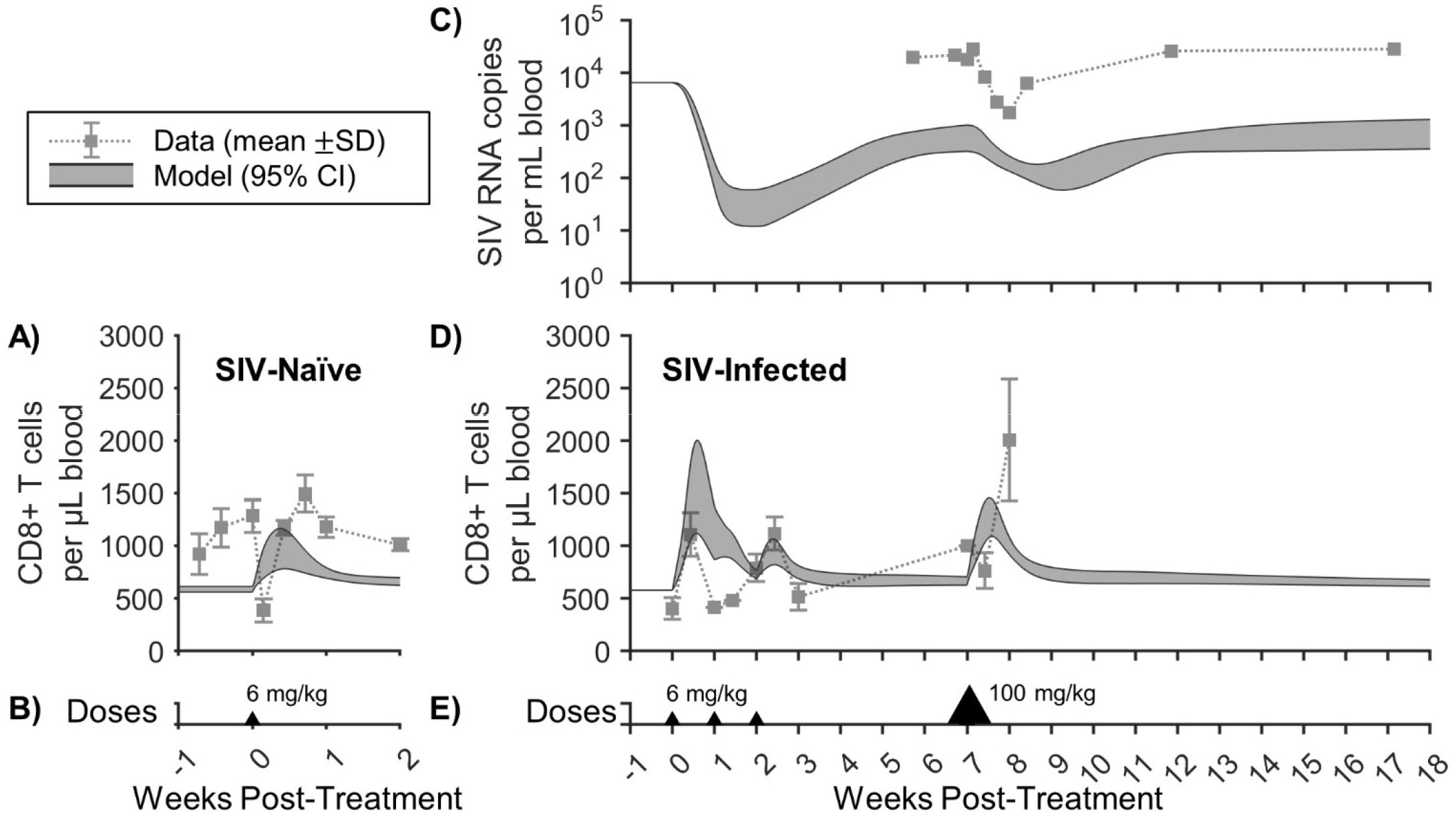
Validation of model with other non-human primate cohorts. Model predictions are compared to a different SIV NHP cohort [21]. In panel A, SIV-naïve NHPs (n=4) are given a 6 mg/kg intravenous dose of N-803 at week 0. In panel D, SIV-infected NHPs (n=4) are given three 6 mg/kg doses spaced one week apart, followed by a 100 mg/kg dose 5 weeks later. Peripheral blood CD8^+^ T cell counts are shown in both panels (mean and standard deviation) and compared to mathematical model predictions (Bayesian 95% credible interval). Panel C shows the plasma viral load after the 100 mg/kg dose (one data subject compared to model prediction). NHP data was obtained from published figures using Engage Digitizer software. Panels (B,E) show timing and size of intravenous doses of N-803.

## References

1. Chauhan P, Nair A, Patidar A, Dandapat J, Sarkar A, Saha B. A primer on cytokines. Cytokine. 2021;145:155458. Epub 2021/02/15. doi: 10.1016/j.cyto.2021.155458. PubMed PMID: 33581983.

2. Lin JX, Leonard WJ. The Common Cytokine Receptor? Chain Family of Cytokines. Cold Spring Harbor perspectives in biology. 2018;10(9). Epub 2017/10/19. doi: 10.1101/cshperspect.a028449. PubMed PMID: 29038115; PubMed Central PMCID: PMCPMC6120701.

3. Conlon KC, Miljkovic MD, Waldmann TA. Cytokines in the Treatment of Cancer. Journal of interferon & cytokine research : the official journal of the International Society for Interferon and Cytokine Research. 2019;39(1):6–21. Epub 2018/06/12. doi: 10.1089/jir.2018.0019. PubMed PMID: 29889594; PubMed Central PMCID: PMCPMC6350412.

4. Hoang TN, Paiardini M. Role of cytokine agonists and immune checkpoint inhibitors toward HIV remission. Current opinion in HIV and AIDS. 2019;14(2):121–8. Epub 2018/12/27. doi: 10.1097/coh.0000000000000528. PubMed PMID: 30585798; PubMed Central PMCID: PMCPMC6469389.

5. Harwood O, O’Connor S. Therapeutic Potential of IL-15 and N-803 in HIV/SIV Infection. Viruses. 2021;13(9). Epub 2021/09/29. doi: 10.3390/v13091750. PubMed PMID: 34578331; PubMed Central PMCID: PMCPMC8473246.

6. Robinson TO, Schluns KS. The potential and promise of IL-15 in immuno-oncogenic therapies. Immunology letters. 2017;190:159–68. Epub 2017/08/22. doi: 10.1016/j.imlet.2017.08.010. PubMed PMID: 28823521; PubMed Central PMCID: PMCPMC5774016.

7. Patidar M, Yadav N, Dalai SK. Interleukin 15: A key cytokine for immunotherapy. Cytokine & growth factor reviews. 2016;31:49–59. Epub 2016/06/22. doi: 10.1016/j.cytogfr.2016.06.001. PubMed PMID: 27325459.

8. Ellis-Connell AL, Balgeman AJ, Zarbock KR, Barry G, Weiler A, Egan JO, et al. ALT-803 Transiently Reduces Simian Immunodeficiency Virus Replication in the Absence of Antiretroviral Treatment. Journal of virology. 2018;92(3). Epub 2017/11/10. doi: 10.1128/jvi.01748-17. PubMed PMID: 29118125; PubMed Central PMCID: PMCPMC5774892.

9. Ellis-Connell AL, Balgeman AJ, Harwood OE, Moriarty RV, Safrit JT, Weiler AM, et al. Control of Simian Immunodeficiency Virus Infection in Prophylactically Vaccinated, Antiretroviral Treatment-Naive Macaques Is Required for the Most Efficacious CD8 T Cell Response during Treatment with the Interleukin-15 Superagonist N-803. Journal of virology. 2022;96(20):e0118522. Epub 2022/10/04. doi: 10.1128/jvi.01185-22. PubMed PMID: 36190241; PubMed Central PMCID: PMCPMC9599604.

10. Webb GM, Li S, Mwakalundwa G, Folkvord JM, Greene JM, Reed JS, et al. The human IL-15 superagonist ALT-803 directs SIV-specific CD8(+) T cells into B-cell follicles. Blood advances. 2018;2(2):76–84. Epub 2018/01/25. doi: 10.1182/bloodadvances.2017012971. PubMed PMID: 29365313; PubMed Central PMCID: PMCPMC5787870 Corporation. The remaining authors declare no competing financial interests.

11. Hatziioannou T, Evans DT. Animal models for HIV/AIDS research. Nature reviews Microbiology. 2012;10(12):852–67. Epub 2012/11/17. doi: 10.1038/nrmicro2911. PubMed PMID: 23154262; PubMed Central PMCID: PMCPMC4334372.

12. Wrangle JM, Velcheti V, Patel MR, Garrett-Mayer E, Hill EG, Ravenel JG, et al. ALT-803, an IL-15 superagonist, in combination with nivolumab in patients with metastatic non-small cell lung cancer: a non-randomised, open-label, phase 1b trial. The Lancet Oncology. 2018;19(5):694–704. Epub 2018/04/10. doi: 10.1016/s1470-2045(18)30148-7. PubMed PMID: 29628312; PubMed Central PMCID: PMCPMC6089612.

13. Margolin K, Morishima C, Velcheti V, Miller JS, Lee SM, Silk AW, et al. Phase I Trial of ALT-803, A Novel Recombinant IL15 Complex, in Patients with Advanced Solid Tumors. Clinical cancer research : an official journal of the American Association for Cancer Research. 2018. Epub 2018/07/27. doi: 10.1158/1078-0432.Ccr-18-0945. PubMed PMID: 30045932.

14. Miller JS, Davis ZB, Helgeson E, Reilly C, Thorkelson A, Anderson J, et al. Safety and virologic impact of the IL-15 superagonist N-803 in people living with HIV: a phase 1 trial. Nature medicine. 2022;28(2):392–400. Epub 2022/02/02. doi: 10.1038/s41591-021-01651-9. PubMed PMID: 35102335.

15. Rhode PR, Egan JO, Xu W, Hong H, Webb GM, Chen X, et al. Comparison of the Superagonist Complex, ALT-803, to IL15 as Cancer Immunotherapeutics in Animal Models. Cancer immunology research. 2016;4(1):49–60. Epub 2015/10/30. doi: 10.1158/2326-6066.cir-15-0093-t. PubMed PMID: 26511282; PubMed Central PMCID: PMCPMC4703482.

16. Padmanabhan P, Dixit NM. Models of Viral Population Dynamics. Current topics in microbiology and immunology. 2016;392:277–302. Epub 2015/07/16. doi: 10.1007/82_2015_458. PubMed PMID: 26174625.

17. Perelson AS, Ribeiro RM. Modeling the within-host dynamics of HIV infection. BMC biology. 2013;11:96. Epub 2013/09/12. doi: 10.1186/1741-7007-11-96. PubMed PMID: 24020860; PubMed Central PMCID: PMCPMC3765939.

18. Simonov M, Rawlings RA, Comment N, Reed SE, Shi X, Nelson PW. Modeling adaptive regulatory T-cell dynamics during early HIV infection. PloS one. 2012;7(4):e33924. Epub 2012/04/27. doi: 10.1371/journal.pone.0033924. PubMed PMID: 22536321; PubMed Central PMCID: PMCPMC3334930.

19. Zheltkova V, Argilaguet J, Peligero C, Bocharov G, Meyerhans A. Prediction of PD-L1 inhibition effects for HIV-infected individuals. PLoS computational biology. 2019;15(11):e1007401. Epub 2019/11/07. doi: 10.1371/journal.pcbi.1007401. PubMed PMID: 31693657; PubMed Central PMCID: PMCPMC6834253.

20. McCann CD, van Dorp CH, Danesh A, Ward AR, Dilling TR, Mota TM, et al. A participant-derived xenograft model of HIV enables long-term evaluation of autologous immunotherapies. The Journal of experimental medicine. 2021;218(7). Epub 2021/05/15. doi: 10.1084/jem.20201908. PubMed PMID: 33988715; PubMed Central PMCID: PMCPMC8129803 “other” from Bristol Myers Squibb outside the submitted work; in addition, D.S. Jones had a patent to PCT/US2019/061837 pending and a patent to PCT/US2017/037249 pending. T.L. Andresen reported being a co-founder of Repertoire Immune Medicine. C.M. Bollard reported “other” from Mana Therapeutics outside the submitted work; in addition, C.M. Bollard had a patent number 9,885,021 licensed to Mana Therapeutics. C.M. Bollard also reported being on the board of directors of Cabaletta Bio, being a co-founder of Mana Therapeutics and Catamaran Bio, and having stock ownership in Repertoire Immune Medicines and Neximmune. D.J. Irvine reported “other” from Repertoire Immune Medicines during the conduct of the study; in addition, D.J. Irvine had a patent to Cell Surface Coupling of Nanoparticles with royalties paid for Repertoire Immune Medicine. No other disclosures were reported.

21. Cody JW, Ellis-Connell AL, O’Connor SL, Pienaar E. Mathematical modeling of N-803 treatment in SIV-infected non-human primates. PLoS computational biology. 2021;17(7):e1009204. Epub 2021/07/29. doi: 10.1371/journal.pcbi.1009204. PubMed PMID: 34319980; PubMed Central PMCID: PMCPMC8351941.

22. Day CL, Kaufmann DE, Kiepiela P, Brown JA, Moodley ES, Reddy S, et al. PD-1 expression on HIV-specific T cells is associated with T-cell exhaustion and disease progression. Nature. 2006;443(7109):350–4. Epub 2006/08/22. doi: 10.1038/nature05115. PubMed PMID: 16921384.

23. Trautmann L, Janbazian L, Chomont N, Said EA, Gimmig S, Bessette B, et al. Upregulation of PD-1 expression on HIV-specific CD8+ T cells leads to reversible immune dysfunction. Nature medicine. 2006;12(10):1198–202. Epub 2006/08/19. doi: 10.1038/nm1482. PubMed PMID: 16917489.

24. Schulze Zur Wiesch J, Thomssen A, Hartjen P, Tóth I, Lehmann C, Meyer-Olson D, et al. Comprehensive analysis of frequency and phenotype of T regulatory cells in HIV infection: CD39 expression of FoxP3+ T regulatory cells correlates with progressive disease. Journal of virology. 2011;85(3):1287–97. Epub 2010/11/05. doi: 10.1128/jvi.01758-10. PubMed PMID: 21047964; PubMed Central PMCID: PMCPMC3020516.

25. Andersson J, Boasso A, Nilsson J, Zhang R, Shire NJ, Lindback S, et al. The prevalence of regulatory T cells in lymphoid tissue is correlated with viral load in HIV-infected patients. Journal of immunology (Baltimore, Md : 1950). 2005;174(6):3143–7. Epub 2005/03/08. doi: 10.4049/jimmunol.174.6.3143. PubMed PMID: 15749840.

26. Fenwick C, Joo V, Jacquier P, Noto A, Banga R, Perreau M, et al. T-cell exhaustion in HIV infection. Immunological reviews. 2019;292(1):149–63. Epub 2019/12/29. doi: 10.1111/imr.12823. PubMed PMID: 31883174; PubMed Central PMCID: PMCPMC7003858.

27. Kleinman AJ, Sivanandham R, Pandrea I, Chougnet CA, Apetrei C. Regulatory T Cells As Potential Targets for HIV Cure Research. Frontiers in immunology. 2018;9:734. Epub 2018/05/01. doi: 10.3389/fimmu.2018.00734. PubMed PMID: 29706961; PubMed Central PMCID: PMCPMC5908895.

28. Penaloza-MacMaster P. CD8 T-cell regulation by T regulatory cells and the programmed cell death protein 1 pathway. Immunology. 2017;151(2):146–53. Epub 2017/04/05. doi: 10.1111/imm.12739. PubMed PMID: 28375543; PubMed Central PMCID: PMCPMC5418463.

29. Waldmann TA, Dubois S, Miljkovic MD, Conlon KC. IL-15 in the Combination Immunotherapy of Cancer. Frontiers in immunology. 2020;11:868. Epub 2020/06/09. doi: 10.3389/fimmu.2020.00868. PubMed PMID: 32508818; PubMed Central PMCID: PMCPMC7248178.

30. Jochems C, Tritsch SR, Knudson KM, Gameiro SR, Rumfield CS, Pellom ST, et al. The multi-functionality of N-809, a novel fusion protein encompassing anti-PD-L1 and the IL-15 superagonist fusion complex. Oncoimmunology. 2019;8(2):e1532764. Epub 2019/02/05. doi: 10.1080/2162402x.2018.1532764. PubMed PMID: 30713787; PubMed Central PMCID: PMCPMC6343815.

31. Knudson KM, Hicks KC, Ozawa Y, Schlom J, Gameiro SR. Functional and mechanistic advantage of the use of a bifunctional anti-PD-L1/IL-15 superagonist. Journal for immunotherapy of cancer. 2020;8(1). Epub 2020/04/19. doi: 10.1136/jitc-2019-000493. PubMed PMID: 32303618; PubMed Central PMCID: PMCPMC7204804.

32. Han KP, Zhu X, Liu B, Jeng E, Kong L, Yovandich JL, et al. IL-15:IL-15 receptor alpha superagonist complex: high-level co-expression in recombinant mammalian cells, purification and characterization. Cytokine. 2011;56(3):804–10. Epub 2011/10/25. doi: 10.1016/j.cyto.2011.09.028. PubMed PMID: 22019703; PubMed Central PMCID: PMCPMC3221918.

33. Betts MR, Ambrozak DR, Douek DC, Bonhoeffer S, Brenchley JM, Casazza JP, et al. Analysis of total human immunodeficiency virus (HIV)-specific CD4(+) and CD8(+) T-cell responses: relationship to viral load in untreated HIV infection. Journal of virology. 2001;75(24):11983–91. Epub 2001/11/17. doi: 10.1128/jvi.75.24.11983-11991.2001. PubMed PMID: 11711588; PubMed Central PMCID: PMCPMC116093.

34. Migueles SA, Connors M. Frequency and function of HIV-specific CD8(+) T cells. Immunology letters. 2001;79(1-2):141–50. Epub 2001/10/12. doi: 10.1016/s0165-2478(01)00276-0. PubMed PMID: 11595301.

35. Gea-Banacloche JC, Migueles SA, Martino L, Shupert WL, McNeil AC, Sabbaghian MS, et al. Maintenance of large numbers of virus-specific CD8+ T cells in HIV-infected progressors and long-term nonprogressors. Journal of immunology (Baltimore, Md : 1950). 2000;165(2):1082–92. Epub 2000/07/06. doi: 10.4049/jimmunol.165.2.1082. PubMed PMID: 10878387.

36. Gadhamsetty S, Beltman JB, de Boer RJ. What do mathematical models tell us about killing rates during HIV-1 infection? Immunology letters. 2015;168(1):1–6. Epub 2015/08/19. doi: 10.1016/j.imlet.2015.07.009. PubMed PMID: 26279491.

37. Kim J, Chang DY, Lee HW, Lee H, Kim JH, Sung PS, et al. Innate-like Cytotoxic Function of Bystander-Activated CD8(+) T Cells Is Associated with Liver Injury in Acute Hepatitis A. Immunity. 2018;48(1):161-73.e5. Epub 2018/01/07. doi: 10.1016/j.immuni.2017.11.025. PubMed PMID: 29305140.

38. Jin X, Bauer DE, Tuttleton SE, Lewin S, Gettie A, Blanchard J, et al. Dramatic rise in plasma viremia after CD8(+) T cell depletion in simian immunodeficiency virus-infected macaques. The Journal of experimental medicine. 1999;189(6):991–8. Epub 1999/03/17. doi: 10.1084/jem.189.6.991. PubMed PMID: 10075982; PubMed Central PMCID: PMCPMC2193038.

39. Choi EI, Reimann KA, Letvin NL. In vivo natural killer cell depletion during primary simian immunodeficiency virus infection in rhesus monkeys. Journal of virology. 2008;82(13):6758–61. Epub 2008/04/25. doi: 10.1128/jvi.02277-07. PubMed PMID: 18434394; PubMed Central PMCID: PMCPMC2447079.

40. Wick WD, Yang OO. Biologically-directed modeling reflects cytolytic clearance of SIV-infected cells in vivo in macaques. PloS one. 2012;7(9):e44778. Epub 2012/10/03. doi: 10.1371/journal.pone.0044778. PubMed PMID: 23028619; PubMed Central PMCID: PMCPMC3441463.

41. Jones LE, Perelson AS. Transient viremia, plasma viral load, and reservoir replenishment in HIV-infected patients on antiretroviral therapy. Journal of acquired immune deficiency syndromes (1999). 2007;45(5):483–93. Epub 2007/05/15. doi: 10.1097/QAI.0b013e3180654836. PubMed PMID: 17496565; PubMed Central PMCID: PMCPMC2584971.

42. De Boer RJ, Mohri H, Ho DD, Perelson AS. Turnover rates of B cells, T cells, and NK cells in simian immunodeficiency virus-infected and uninfected rhesus macaques. Journal of immunology (Baltimore, Md : 1950). 2003;170(5):2479–87. Epub 2003/02/21. doi: 10.4049/jimmunol.170.5.2479. PubMed PMID: 12594273.

43. Romee R, Cooley S, Berrien-Elliott MM, Westervelt P, Verneris MR, Wagner JE, et al. First-in-human phase 1 clinical study of the IL-15 superagonist complex ALT-803 to treat relapse after transplantation. Blood. 2018;131(23):2515–27. Epub 2018/02/22. doi: 10.1182/blood-2017-12-823757. PubMed PMID: 29463563; PubMed Central PMCID: PMCPMC5992862 support from Altor BioScience, a Nantworks company, but have no financial benefit from the outcome of this trial. J.O.E., E.K.J., A.R., and H.C.W. are employees of Altor BioScience and declare direct financial conflicts. To manage these conflicts, UMN and WUSM investigators led this trial, were sponsors of the IND, managed all the data in the study, and had final responsibility for the manuscript. The study protocol was an investigator-initiated clinical trial. UMN and WUSM investigators performed the clinical trial including data collection, analysis, and interpretation. Altor BioScience performed ALT-803 and cytokine measurements and immunogenicity testing on coded samples. The remaining correlative assays and all statistical analyses were performed by UMN and WUSM. The remaining authors declare no competing financial interests.

44. Davenport MP, Ribeiro RM, Perelson AS. Kinetics of virus-specific CD8+ T cells and the control of human immunodeficiency virus infection. Journal of virology. 2004;78(18):10096–103. Epub 2004/08/28. doi: 10.1128/jvi.78.18.10096-10103.2004. PubMed PMID: 15331742; PubMed Central PMCID: PMCPMC515020.

45. Cardozo EF, Andrade A, Mellors JW, Kuritzkes DR, Perelson AS, Ribeiro RM. Treatment with integrase inhibitor suggests a new interpretation of HIV RNA decay curves that reveals a subset of cells with slow integration. PLoS pathogens. 2017;13(7):e1006478. Epub 2017/07/06. doi: 10.1371/journal.ppat.1006478. PubMed PMID: 28678879; PubMed Central PMCID: PMCPMC5513547.

46. Conway JM, Perelson AS. Residual Viremia in Treated HIV+ Individuals. PLoS computational biology. 2016;12(1):e1004677. Epub 2016/01/07. doi: 10.1371/journal.pcbi.1004677. PubMed PMID: 26735135; PubMed Central PMCID: PMCPMC4703306.

47. Chun TW, Carruth L, Finzi D, Shen X, DiGiuseppe JA, Taylor H, et al. Quantification of latent tissue reservoirs and total body viral load in HIV-1 infection. Nature. 1997;387(6629):183–8. Epub 1997/05/08. doi: 10.1038/387183a0. PubMed PMID: 9144289.

48. Choo DK, Murali-Krishna K, Anita R, Ahmed R. Homeostatic turnover of virus-specific memory CD8 T cells occurs stochastically and is independent of CD4 T cell help. Journal of immunology (Baltimore, Md : 1950). 2010;185(6):3436–44. Epub 2010/08/25. doi: 10.4049/jimmunol.1001421. PubMed PMID: 20733203.

49. Hataye J, Moon JJ, Khoruts A, Reilly C, Jenkins MK. Naive and memory CD4+ T cell survival controlled by clonal abundance. Science (New York, NY). 2006;312(5770):114–6. Epub 2006/03/04. doi: 10.1126/science.1124228. PubMed PMID: 16513943.

50. Freitas AA, Rocha B. Population biology of lymphocytes: the flight for survival. Annual review of immunology. 2000;18:83–111. Epub 2000/06/03. doi: 10.1146/annurev.immunol.18.1.83. PubMed PMID: 10837053.

51. Younes SA, Freeman ML, Mudd JC, Shive CL, Reynaldi A, Panigrahi S, et al. IL-15 promotes activation and expansion of CD8+ T cells in HIV-1 infection. The Journal of clinical investigation. 2016;126(7):2745–56. Epub 2016/06/21. doi: 10.1172/jci85996. PubMed PMID: 27322062; PubMed Central PMCID: PMCPMC4922693.

52. Bastidas S, Graw F, Smith MZ, Kuster H, Günthard HF, Oxenius A. CD8+ T cells are activated in an antigen-independent manner in HIV-infected individuals. Journal of immunology (Baltimore, Md : 1950). 2014;192(4):1732–44. Epub 2014/01/22. doi: 10.4049/jimmunol.1302027. PubMed PMID: 24446519.

53. Doisne JM, Urrutia A, Lacabaratz-Porret C, Goujard C, Meyer L, Chaix ML, et al. CD8+ T cells specific for EBV, cytomegalovirus, and influenza virus are activated during primary HIV infection. Journal of immunology (Baltimore, Md : 1950). 2004;173(4):2410–8. Epub 2004/08/06. doi: 10.4049/jimmunol.173.4.2410. PubMed PMID: 15294954.

54. Jambhekar SS, Breen PJ. Extravascular routes of drug administration. Basic Pharmacokinetics. 2 ed: Pharmaceutical Press; 2012. p. 105–26.

55. Richer MJ, Pewe LL, Hancox LS, Hartwig SM, Varga SM, Harty JT. Inflammatory IL-15 is required for optimal memory T cell responses. The Journal of clinical investigation. 2015;125(9):3477–90. Epub 2015/08/05. doi: 10.1172/jci81261. PubMed PMID: 26241055; PubMed Central PMCID: PMCPMC4588296.

56. Vang KB, Yang J, Mahmud SA, Burchill MA, Vegoe AL, Farrar MA. IL-2, -7, and -15, but not thymic stromal lymphopoeitin, redundantly govern CD4+Foxp3+ regulatory T cell development. Journal of immunology (Baltimore, Md : 1950). 2008;181(5):3285–90. Epub 2008/08/21. doi: 10.4049/jimmunol.181.5.3285. PubMed PMID: 18714000; PubMed Central PMCID: PMCPMC2810104.

57. Kinter AL, Godbout EJ, McNally JP, Sereti I, Roby GA, O’Shea MA, et al. The common gamma-chain cytokines IL-2, IL-7, IL-15, and IL-21 induce the expression of programmed death-1 and its ligands. Journal of immunology (Baltimore, Md : 1950). 2008;181(10):6738–46. Epub 2008/11/05. doi: 10.4049/jimmunol.181.10.6738. PubMed PMID: 18981091.

58. Burrack KS, Huggins MA, Taras E, Dougherty P, Henzler CM, Yang R, et al. Interleukin-15 Complex Treatment Protects Mice from Cerebral Malaria by Inducing Interleukin-10-Producing Natural Killer Cells. Immunity. 2018;48(4):760-72.e4. Epub 2018/04/08. doi: 10.1016/j.immuni.2018.03.012. PubMed PMID: 29625893; PubMed Central PMCID: PMCPMC5906161.

59. Kim PS, Kwilas AR, Xu W, Alter S, Jeng EK, Wong HC, et al. IL-15 superagonist/IL-15RalphaSushi-Fc fusion complex (IL-15SA/IL-15RalphaSu-Fc; ALT-803) markedly enhances specific subpopulations of NK and memory CD8+ T cells, and mediates potent anti-tumor activity against murine breast and colon carcinomas. Oncotarget. 2016;7(13):16130–45. Epub 2016/02/26. doi: 10.18632/oncotarget.7470. PubMed PMID: 26910920; PubMed Central PMCID: PMCPMC4941302.

60. Martins MA, Tully DC, Cruz MA, Power KA, Veloso de Santana MG, Bean DJ, et al. Vaccine-Induced Simian Immunodeficiency Virus-Specific CD8+ T-Cell Responses Focused on a Single Nef Epitope Select for Escape Variants Shortly after Infection. Journal of virology. 2015;89(21):10802–20. Epub 2015/08/21. doi: 10.1128/jvi.01440-15. PubMed PMID: 26292326; PubMed Central PMCID: PMCPMC4621113.

61. Loffredo JT, Maxwell J, Qi Y, Glidden CE, Borchardt GJ, Soma T, et al. Mamu-B*08-positive macaques control simian immunodeficiency virus replication. Journal of virology. 2007;81(16):8827–32. Epub 2007/06/01. doi: 10.1128/jvi.00895-07. PubMed PMID: 17537848; PubMed Central PMCID: PMCPMC1951344.

62. Pinkevych M, Fennessey CM, Cromer D, Tolstrup M, Sogaard OS, Rasmussen TA, et al. Estimating Initial Viral Levels during Simian Immunodeficiency Virus/Human Immunodeficiency Virus Reactivation from Latency. Journal of virology. 2018;92(2). Epub 2017/11/10. doi: 10.1128/jvi.01667-17. PubMed PMID: 29118123; PubMed Central PMCID: PMCPMC5752936.

63. McKay MD, Beckman RJ, Conover WJ. Comparison of Three Methods for Selecting Values of Input Variables in the Analysis of Output from a Computer Code. Technometrics. 1979;21(2):239–45. doi: 10.1080/00401706.1979.10489755.

64. Byrd RH, Gilbert JC, Nocedal J. A trust region method based on interior point techniques for nonlinear programming. Mathematical Programming. 2000;89(1):149–85. doi: 10.1007/PL00011391. PubMed PMID: NA.

65. Vousden WD, Farr WM, Mandel I. Dynamic temperature selection for parallel tempering in Markov chain Monte Carlo simulations. Monthly Notices of the Royal Astronomical Society. 2015;455(2):1919–37. doi: 10.1093/mnras/stv2422. PubMed PMID: NA.

66. Weninger W, Crowley MA, Manjunath N, von Andrian UH. Migratory properties of naive, effector, and memory CD8(+) T cells. The Journal of experimental medicine. 2001;194(7):953–66. Epub 2001/10/03. doi: 10.1084/jem.194.7.953. PubMed PMID: 11581317; PubMed Central PMCID: PMCPMC2193483.

67. Sowell RT, Goldufsky JW, Rogozinska M, Quiles Z, Cao Y, Castillo EF, et al. IL-15 Complexes Induce Migration of Resting Memory CD8 T Cells into Mucosal Tissues. Journal of immunology (Baltimore, Md : 1950). 2017;199(7):2536–46. Epub 2017/08/18. doi: 10.4049/jimmunol.1501638. PubMed PMID: 28814601; PubMed Central PMCID: PMCPMC5605445.

68. Voskoboinik I, Whisstock JC, Trapani JA. Perforin and granzymes: function, dysfunction and human pathology. Nature reviews Immunology. 2015;15(6):388–400. Epub 2015/05/23. doi: 10.1038/nri3839. PubMed PMID: 25998963.

69. Akhmetzyanova I, Drabczyk M, Neff CP, Gibbert K, Dietze KK, Werner T, et al. PD-L1 Expression on Retrovirus-Infected Cells Mediates Immune Escape from CD8+ T Cell Killing. PLoS pathogens. 2015;11(10):e1005224. Epub 2015/10/21. doi: 10.1371/journal.ppat.1005224. PubMed PMID: 26484769; PubMed Central PMCID: PMCPMC4617866.

70. Gonzalo-Gil E, Ikediobi U, Sutton RE. Mechanisms of Virologic Control and Clinical Characteristics of HIV+ Elite/Viremic Controllers. The Yale journal of biology and medicine. 2017;90(2):245–59. Epub 2017/06/29. PubMed PMID: 28656011; PubMed Central PMCID: PMCPMC5482301.

71. Mastrangelo A, Banga R, Perreau M. Elite and posttreatment controllers, two facets of HIV control. Current opinion in HIV and AIDS. 2022;17(5):325–32. Epub 2022/08/09. doi: 10.1097/coh.0000000000000751. PubMed PMID: 35938466.

72. Penaloza-MacMaster P, Kamphorst AO, Wieland A, Araki K, Iyer SS, West EE, et al. Interplay between regulatory T cells and PD-1 in modulating T cell exhaustion and viral control during chronic LCMV infection. The Journal of experimental medicine. 2014;211(9):1905–18. Epub 2014/08/13. doi: 10.1084/jem.20132577. PubMed PMID: 25113973; PubMed Central PMCID: PMCPMC4144726.

73. Buchbinder EI, Desai A. CTLA-4 and PD-1 Pathways: Similarities, Differences, and Implications of Their Inhibition. American journal of clinical oncology. 2016;39(1):98–106. Epub 2015/11/13. doi: 10.1097/coc.0000000000000239. PubMed PMID: 26558876; PubMed Central PMCID: PMCPMC4892769 spouse previously employed by Merck. A.D. declares no conflicts of interest.

74. Garner H, de Visser KE. Immune crosstalk in cancer progression and metastatic spread: a complex conversation. Nature reviews Immunology. 2020;20(8):483–97. Epub 2020/02/07. doi: 10.1038/s41577-019-0271-z. PubMed PMID: 32024984.

75. Yu P, Steel JC, Zhang M, Morris JC, Waitz R, Fasso M, et al. Simultaneous inhibition of two regulatory T-cell subsets enhanced Interleukin-15 efficacy in a prostate tumor model. Proceedings of the National Academy of Sciences of the United States of America. 2012;109(16):6187–92. Epub 2012/04/05. doi: 10.1073/pnas.1203479109. PubMed PMID: 22474386; PubMed Central PMCID: PMCPMC3341063.

76. Gabrilovich DI. Myeloid-Derived Suppressor Cells. Cancer immunology research. 2017;5(1):3–8. Epub 2017/01/06. doi: 10.1158/2326-6066.Cir-16-0297. PubMed PMID: 28052991; PubMed Central PMCID: PMCPMC5426480.

77. Highfill SL, Cui Y, Giles AJ, Smith JP, Zhang H, Morse E, et al. Disruption of CXCR2-mediated MDSC tumor trafficking enhances anti-PD1 efficacy. Science translational medicine. 2014;6(237):237ra67. Epub 2014/05/23. doi: 10.1126/scitranslmed.3007974. PubMed PMID: 24848257; PubMed Central PMCID: PMCPMC6980372.

78. Meyer C, Cagnon L, Costa-Nunes CM, Baumgaertner P, Montandon N, Leyvraz L, et al. Frequencies of circulating MDSC correlate with clinical outcome of melanoma patients treated with ipilimumab. Cancer immunology, immunotherapy : CII. 2014;63(3):247–57. Epub 2013/12/21. doi: 10.1007/s00262-013-1508-5. PubMed PMID: 24357148.

79. Fabian KP, Padget MR, Fujii R, Schlom J, Hodge JW. Differential combination immunotherapy requirements for inflamed (warm) tumors versus T cell excluded (cool) tumors: engage, expand, enable, and evolve. Journal for immunotherapy of cancer. 2021;9(2). Epub 2021/02/20. doi: 10.1136/jitc-2020-001691. PubMed PMID: 33602696; PubMed Central PMCID: PMCPMC7896589.

80. Drusbosky L, Nangia C, Nguyen A, Szeto C, Newton Y, Spilman P, et al. Complete response to avelumab and IL-15 superagonist N-803 with Abraxane in Merkel cell carcinoma: a case study. Journal for immunotherapy of cancer. 2020;8(2). Epub 2020/09/12. doi: 10.1136/jitc-2020-001098. PubMed PMID: 32913030; PubMed Central PMCID: PMCPMC7484858.

81. Hodge JW, Garnett CT, Farsaci B, Palena C, Tsang KY, Ferrone S, et al. Chemotherapy-induced immunogenic modulation of tumor cells enhances killing by cytotoxic T lymphocytes and is distinct from immunogenic cell death. International journal of cancer. 2013;133(3):624–36. Epub 2013/02/01. doi: 10.1002/ijc.28070. PubMed PMID: 23364915; PubMed Central PMCID: PMCPMC3663913.

82. Albeituni SH, Ding C, Yan J. Hampering immune suppressors: therapeutic targeting of myeloid-derived suppressor cells in cancer. Cancer journal (Sudbury, Mass). 2013;19(6):490–501. Epub 2013/11/26. doi: 10.1097/ppo.0000000000000006. PubMed PMID: 24270348; PubMed Central PMCID: PMCPMC3902636.

83. Gaither KA, Little AA, McBride AA, Garcia SR, Brar KK, Zhu Z, et al. The immunomodulatory, antitumor and antimetastatic responses of melanoma-bearing normal and alcoholic mice to sunitinib and ALT-803: a combinatorial treatment approach. Cancer immunology, immunotherapy : CII. 2016;65(9):1123–34. Epub 2016/08/03. doi: 10.1007/s00262-016-1876-8. PubMed PMID: 27481107.

84. Streeck H, Brumme ZL, Anastario M, Cohen KW, Jolin JS, Meier A, et al. Antigen load and viral sequence diversification determine the functional profile of HIV-1-specific CD8+ T cells. PLoS medicine. 2008;5(5):e100. Epub 2008/05/09. doi: 10.1371/journal.pmed.0050100. PubMed PMID: 18462013; PubMed Central PMCID: PMCPMC2365971.

85. Pallikkuth S, Fischl MA, Pahwa S. Combination antiretroviral therapy with raltegravir leads to rapid immunologic reconstitution in treatment-naive patients with chronic HIV infection. The Journal of infectious diseases. 2013;208(10):1613–23. Epub 2013/08/08. doi: 10.1093/infdis/jit387. PubMed PMID: 23922374; PubMed Central PMCID: PMCPMC3805240.

86. Presicce P, Orsborn K, King E, Pratt J, Fichtenbaum CJ, Chougnet CA. Frequency of circulating regulatory T cells increases during chronic HIV infection and is largely controlled by highly active antiretroviral therapy. PloS one. 2011;6(12):e28118. Epub 2011/12/14. doi: 10.1371/journal.pone.0028118. PubMed PMID: 22162758; PubMed Central PMCID: PMCPMC3230597 Gilead Sciences. This does not alter the authors’ adherence to all PLoS ONE policies on sharing data and materials.

87. Epple HJ, Loddenkemper C, Kunkel D, Tröger H, Maul J, Moos V, et al. Mucosal but not peripheral FOXP3+ regulatory T cells are highly increased in untreated HIV infection and normalize after suppressive HAART. Blood. 2006;108(9):3072–8. Epub 2006/05/27. doi: 10.1182/blood-2006-04-016923. PubMed PMID: 16728694.

88. McBrien JB, Mavigner M, Franchitti L, Smith SA, White E, Tharp GK, et al. Robust and persistent reactivation of SIV and HIV by N-803 and depletion of CD8(+) cells. Nature. 2020;578(7793):154–9. Epub 2020/01/24. doi: 10.1038/s41586-020-1946-0. PubMed PMID: 31969705.

89. Briesacher BA, Andrade SE, Harrold LR, Fouayzi H, Yood RA. Adoption of once-monthly oral bisphosphonates and the impact on adherence. The American journal of medicine. 2010;123(3):275–80. Epub 2010/03/03. doi: 10.1016/j.amjmed.2009.05.017. PubMed PMID: 20193837; PubMed Central PMCID: PMCPMC2831769.

90. Iglay K, Cao X, Mavros P, Joshi K, Yu S, Tunceli K. Systematic Literature Review and Meta-analysis of Medication Adherence With Once-weekly Versus Once-daily Therapy. Clin Ther. 2015;37(8):1813-21.e1. Epub 2015/06/29. doi: 10.1016/j.clinthera.2015.05.505. PubMed PMID: 26117406.

91. Jones RB, Mueller S, O’Connor R, Rimpel K, Sloan DD, Karel D, et al. A Subset of Latency-Reversing Agents Expose HIV-Infected Resting CD4+ T-Cells to Recognition by Cytotoxic T-Lymphocytes. PLoS pathogens. 2016;12(4):e1005545. Epub 2016/04/16. doi: 10.1371/journal.ppat.1005545. PubMed PMID: 27082643; PubMed Central PMCID: PMCPMC4833318.

92. Bronnimann MP, Skinner PJ, Connick E. The B-Cell Follicle in HIV Infection: Barrier to a Cure. Frontiers in immunology. 2018;9:20. Epub 2018/02/10. doi: 10.3389/fimmu.2018.00020. PubMed PMID: 29422894; PubMed Central PMCID: PMCPMC5788973.

## References

1. Ellis-Connell AL, Balgeman AJ, Zarbock KR, Barry G, Weiler A, Egan JO, et al. ALT-803 Transiently Reduces Simian Immunodeficiency Virus Replication in the Absence of Antiretroviral Treatment. Journal of virology. 2018;92(3). Epub 2017/11/10. doi: 10.1128/jvi.01748-17. PubMed PMID: 29118125; PubMed Central PMCID: PMCPMC5774892.

2. Ellis-Connell AL, Balgeman AJ, Harwood OE, Moriarty RV, Safrit JT, Weiler AM, et al. Control of Simian Immunodeficiency Virus Infection in Prophylactically Vaccinated, Antiretroviral Treatment-Naive Macaques Is Required for the Most Efficacious CD8 T Cell Response during Treatment with the Interleukin-15 Superagonist N-803. Journal of virology. 2022;96(20):e0118522. Epub 2022/10/04. doi: 10.1128/jvi.01185-22. PubMed PMID: 36190241; PubMed Central PMCID: PMCPMC9599604.

3. Betts MR, Ambrozak DR, Douek DC, Bonhoeffer S, Brenchley JM, Casazza JP, et al. Analysis of total human immunodeficiency virus (HIV)-specific CD4(+) and CD8(+) T-cell responses: relationship to viral load in untreated HIV infection. Journal of virology. 2001;75(24):11983–91. Epub 2001/11/17. doi: 10.1128/jvi.75.24.11983-11991.2001. PubMed PMID: 11711588; PubMed Central PMCID: PMCPMC116093.

4. Migueles SA, Connors M. Frequency and function of HIV-specific CD8(+) T cells. Immunology letters. 2001;79(1-2):141–50. Epub 2001/10/12. doi: 10.1016/s0165-2478(01)00276-0. PubMed PMID: 11595301.

5. Gea-Banacloche JC, Migueles SA, Martino L, Shupert WL, McNeil AC, Sabbaghian MS, et al. Maintenance of large numbers of virus-specific CD8+ T cells in HIV-infected progressors and long-term nonprogressors. Journal of immunology (Baltimore, Md : 1950). 2000;165(2):1082–92. Epub 2000/07/06. doi: 10.4049/jimmunol.165.2.1082. PubMed PMID: 10878387.

6. Han KP, Zhu X, Liu B, Jeng E, Kong L, Yovandich JL, et al. IL-15:IL-15 receptor alpha superagonist complex: high-level co-expression in recombinant mammalian cells, purification and characterization. Cytokine. 2011;56(3):804–10. Epub 2011/10/25. doi: 10.1016/j.cyto.2011.09.028. PubMed PMID: 22019703; PubMed Central PMCID: PMCPMC3221918.

7. Gadhamsetty S, Beltman JB, de Boer RJ. What do mathematical models tell us about killing rates during HIV-1 infection? Immunology letters. 2015;168(1):1–6. Epub 2015/08/19. doi: 10.1016/j.imlet.2015.07.009. PubMed PMID: 26279491.

8. Kim J, Chang DY, Lee HW, Lee H, Kim JH, Sung PS, et al. Innate-like Cytotoxic Function of Bystander-Activated CD8(+) T Cells Is Associated with Liver Injury in Acute Hepatitis A. Immunity. 2018;48(1):161-73.e5. Epub 2018/01/07. doi: 10.1016/j.immuni.2017.11.025. PubMed PMID: 29305140.

9. Choi EI, Reimann KA, Letvin NL. In vivo natural killer cell depletion during primary simian immunodeficiency virus infection in rhesus monkeys. Journal of virology. 2008;82(13):6758–61. Epub 2008/04/25. doi: 10.1128/jvi.02277-07. PubMed PMID: 18434394; PubMed Central PMCID: PMCPMC2447079.

10. Jin X, Bauer DE, Tuttleton SE, Lewin S, Gettie A, Blanchard J, et al. Dramatic rise in plasma viremia after CD8(+) T cell depletion in simian immunodeficiency virus-infected macaques. The Journal of experimental medicine. 1999;189(6):991–8. Epub 1999/03/17. doi: 10.1084/jem.189.6.991. PubMed PMID: 10075982; PubMed Central PMCID: PMCPMC2193038.

11. Wick WD, Yang OO. Biologically-directed modeling reflects cytolytic clearance of SIV-infected cells in vivo in macaques. PloS one. 2012;7(9):e44778. Epub 2012/10/03. doi: 10.1371/journal.pone.0044778. PubMed PMID: 23028619; PubMed Central PMCID: PMCPMC3441463.

12. Jones LE, Perelson AS. Transient viremia, plasma viral load, and reservoir replenishment in HIV-infected patients on antiretroviral therapy. Journal of acquired immune deficiency syndromes (1999). 2007;45(5):483–93. Epub 2007/05/15. doi: 10.1097/QAI.0b013e3180654836. PubMed PMID: 17496565; PubMed Central PMCID: PMCPMC2584971.

13. Lalvani A, Brookes R, Hambleton S, Britton WJ, Hill AV, McMichael AJ. Rapid effector function in CD8+ memory T cells. The Journal of experimental medicine. 1997;186(6):859–65. Epub 1997/09/18. doi: 10.1084/jem.186.6.859. PubMed PMID: 9294140; PubMed Central PMCID: PMCPMC2199056.

14. De Boer RJ, Mohri H, Ho DD, Perelson AS. Turnover rates of B cells, T cells, and NK cells in simian immunodeficiency virus-infected and uninfected rhesus macaques. Journal of immunology (Baltimore, Md : 1950). 2003;170(5):2479–87. Epub 2003/02/21. doi: 10.4049/jimmunol.170.5.2479. PubMed PMID: 12594273.

15. Davenport MP, Ribeiro RM, Perelson AS. Kinetics of virus-specific CD8+ T cells and the control of human immunodeficiency virus infection. Journal of virology. 2004;78(18):10096–103. Epub 2004/08/28. doi: 10.1128/jvi.78.18.10096-10103.2004. PubMed PMID: 15331742; PubMed Central PMCID: PMCPMC515020.

16. Cody JW, Ellis-Connell AL, O’Connor SL, Pienaar E. Mathematical modeling of N-803 treatment in SIV-infected non-human primates. PLoS computational biology. 2021;17(7):e1009204. Epub 2021/07/29. doi: 10.1371/journal.pcbi.1009204. PubMed PMID: 34319980; PubMed Central PMCID: PMCPMC8351941.

17. Seber GAF, Wild CJ. Nonlinear Regression. Hoboken, NJ: John Wiley & Sons; 2003.

18. McKay MD, Beckman RJ, Conover WJ. Comparison of Three Methods for Selecting Values of Input Variables in the Analysis of Output from a Computer Code. Technometrics. 1979;21(2):239–45. doi: 10.1080/00401706.1979.10489755.

19. Vousden WD, Farr WM, Mandel I. Dynamic temperature selection for parallel tempering in Markov chain Monte Carlo simulations. Monthly Notices of the Royal Astronomical Society. 2015;455(2):1919–37. doi: 10.1093/mnras/stv2422. PubMed PMID: NA.

20. Stapor P, Weindl D, Ballnus B, Hug S, Loos C, Fiedler A, et al. PESTO: Parameter EStimation TOolbox. Bioinformatics (Oxford, England). 2018;34(4):705–7. Epub 2017/10/27. doi: 10.1093/bioinformatics/btx676. PubMed PMID: 29069312; PubMed Central PMCID: PMCPMC5860618.

21. Webb GM, Li S, Mwakalundwa G, Folkvord JM, Greene JM, Reed JS, et al. The human IL-15 superagonist ALT-803 directs SIV-specific CD8(+) T cells into B-cell follicles. Blood advances. 2018;2(2):76–84. Epub 2018/01/25. doi: 10.1182/bloodadvances.2017012971. PubMed PMID: 29365313; PubMed Central PMCID: PMCPMC5787870 Corporation. The remaining authors declare no competing financial interests.

22. Weninger W, Crowley MA, Manjunath N, von Andrian UH. Migratory properties of naive, effector, and memory CD8(+) T cells. The Journal of experimental medicine. 2001;194(7):953–66. Epub 2001/10/03. doi: 10.1084/jem.194.7.953. PubMed PMID: 11581317; PubMed Central PMCID: PMCPMC2193483.

23. Sowell RT, Goldufsky JW, Rogozinska M, Quiles Z, Cao Y, Castillo EF, et al. IL-15 Complexes Induce Migration of Resting Memory CD8 T Cells into Mucosal Tissues. Journal of immunology (Baltimore, Md : 1950). 2017;199(7):2536–46. Epub 2017/08/18. doi: 10.4049/jimmunol.1501638. PubMed PMID: 28814601; PubMed Central PMCID: PMCPMC5605445.

